# Tau Ablation Rewires Brain Cell Programs in Health and Restores Function in Disease

**DOI:** 10.64898/2026.05.12.724684

**Authors:** Enrique Chimal-Juárez, Henika Patel, Nur Jury-Garfe, Luke C. Dabin, Ruben Vidal, Jungsu Kim, Cristian A. Lasagna-Reeves

## Abstract

Tau is a microtubule-associated protein with diverse roles in the healthy brain but contributes to neurodegenerative disorders when dysregulated. Although tau ablation has shown protective effects in several disease models, how its absence confers this protection remain unclear. Here, we performed and analyzed single-nucleus RNA sequencing on cortices of aged tau knockout (*Mapt^⁻/⁻^*) mice in a wild type background as well as in a vascular amyloid model to evaluate the effect on disease context. Comparisons in a wild type setting revealed that tau ablation induced compensatory remodeling across multiple cell types. Excitatory neurons expanded into a distinct subtype with unique glutamatergic signaling, astrocytes adopted synaptoprotective states, oligodendrocytes upregulated genes supporting connectivity and plasticity, and microglia engaged structural remodeling programs. In contrast, in disease, tau removal not only restored functions disrupted by vascular amyloid pathology, but also generated new phenotypes. Excitatory neurons rewired receptor and postsynaptic signaling, astrocytes and oligodendrocytes recovered wild-type-like gene programs related to neurotransmitter cycling, synaptic support, and myelin integrity, and microglia reprogrammed toward sensing and mounting responses. Together, these findings demonstrate that tau ablation reshapes brain cellular programs in a context-dependent manner, exerting adaptive responses in the otherwise healthy brain while restoring homeostatic functions under vascular amyloid pathology. These results position tau as a key regulator of neuronal-glial network balance and highlight the importance of understanding how tau influences distinct cellular programs within specific disease environments.

## INTRODUCTION

The microtubule-associated protein tau (MAPT) was first identified as a key regulator of microtubule stability and assembly [1]. Tau stabilizes microtubules by interacting with tubulin heterodimers, thereby supporting axonal structure and maintaining neuronal morphology [2–4]. Tau is predominantly expressed in cortical and hippocampal neurons [5–7], although there is also expression in other cell types such as oligodendrocytes and astrocytes [8, 9]. While tau is commonly studied in neurodegenerative disorders, its role in the healthy brain remains an important area of investigation. Genetic deletion studies indicate that endogenous tau participates in a range of neuronal processes, including cytoskeletal regulation, mitochondrial dynamics, synaptic plasticity, network activity [10–22], and stability of DNA and RNA stability [23]. In the adult human brain, splicing of the MAPT gene in exons 2, 3, and 10 generates six isoforms, including variants with three or four microtubule-binding repeats [24, 25]. In addition, tau can also be subjected to a myriad of post translational modifications (PTMs) including phosphorylation [26–29], acetylation [30–32], ubiquitination [33, 34], and methylation [35]. The diversity of tau isoforms and PTMs contribute to its functional diversity by regulating its localization, stability, and interactions [36].

Tau-mediated neurodegeneration arises from multiple pathological mechanisms, including loss of normal function [37–42], gain of toxic function [43–46], aggregation [44, 47–49], and mislocalization [50–52]. A prime example are the neurofibrillary tangles (NFTs) in Alzheimer’s Disease (AD), recognized as a pathological hallmark of the disease [53]. Studies have showed that NFTs are primarily composed of hyperphosphorylated aggregates of tau [54]. In addition to classical tauopathies, tau dysregulation has also been implicated in a range of neurological disorders, including autism spectrum disorder (ASD) [55–58], Parkinson’s Disease (PD) [59–63], Huntington’s Disease (HD) [64–68], and Traumatic Brain Injury (TBI) [69–73].

Given tau’s role in neurodegenerative processes and the therapeutic interest in its modulation [74], growing attention has been directed toward tau ablation models to investigate how the loss of tau influences disease-associated phenotypes across neurological disorders. Interestingly, in multiple mouse disease models, tau ablation has been associated with protective effects. For example, in AD models, the absence of tau has been shown to rescue behavioral impairments and reduce neurotoxicity, despite no significant changes in Aβ levels [75–79]. In models of ASD, tau knockdown has been associated with reduced autistic-like behaviors and decreased seizure susceptibility [56, 80, 81]. Some PD models showed improvements in aberrant network activity upon tau removal [82], while in HD models, ablation led to reduced motor abnormalities and restoration of neurite length [83, 84]. Moreover, we recently demonstrated that tau ablation is beneficial in a mouse model of cerebral amyloid angiopathy (CAA), reducing vascular damage and improving behavioral outcomes [85]. Beyond neurodegenerative and neurodevelopmental disorders, tau ablation has also been shown to improve outcomes in models of neurological injury and vascular stress, such TBI and high-salt diet-induced vascular dysfunction, with associated improvements in cognitive performance [72, 86, 87]. Despite these consistent protective effects, the cellular and molecular mechanisms underlying the protection conferred by tau depletion remain poorly understood. Further work is needed to define tau’s physiological roles and to elucidate how its modulation may be therapeutically leveraged, including its impact on cell type-specific transcriptional programs in disease contexts.

Here, we performed snRNA sequencing on *Mapt^⁻/⁻^* mice and compared them to age-matched wild-type (WT) littermates. Our results revealed genotype-dependent changes across multiple brain cell types under physiological conditions. Neurons and astrocytes showed the most pronounced changes in tau’s absence, including shifts in subtype composition and gene expression related to synaptic and support functions, while oligodendrocytes and microglia exhibited mainly transcriptional adaptations linked to connectivity and remodeling. Overall, these findings suggest cell type-specific compensatory responses rather than widespread cellular reorganization. To investigate the cellular and molecular consequences of tau ablation in a disease context, we analyzed the Tg-FDD mouse, a model for CAA, after *Mapt* ablation (Tg-FDD; *Mapt^⁻/⁻^*). Notably, tau ablation in CAA context was associated with a reduction of key pathological phenotypes. Comparative analysis revealed that Tg-FDD;*Mapt^⁻/⁻^* mice have expansion of a specific excitatory neuron subtype characterized by unique glutamatergic and postsynaptic signaling gene programs, suggesting compensatory synaptic plasticity and rewired excitatory connectivity. Although astrocytes, oligodendrocytes, and microglia maintained stable proportions, they exhibited restoration of gene programs that were disrupted in Tg-FDD mice, indicating glial recovery of functions distinct from those observed in loss of tau in a wild type context.

## MATERIALS AND METHODS

### Transgenic mouse model

Twenty-two-month-old *Mapt*^-/-^ (JAX stock #007251), Tg-FDD;*Mapt*^-/-^, Tg-FDD and wild type (WT) male mice were used for our experiments. Male mice were selected due to the strong phenotype and recovery previously described [85]. The Tg-FDD mice express an FDD-associated human mutant BRI2 transgene that leads to the vascular accumulation of the ADan amyloid [88]. Breeding strategy was followed as previously described [85]. Mice were housed at the Indiana University School of Medicine (IUSM) animal care facility and were maintained according to USDA standards (12-hr light/dark cycle, food and water ad libitum), per the Guide for the Care and Use of Laboratory Animals (National Institutes of Health, Bethesda, MD). Animals were deeply anesthetized and euthanized according to the IUSM Institutional Animal Care and Use Committee-approved procedures. After sacrifice, brain tissue was collected and stored at-80 °C [85].

### Singulator S100 nuclei isolation

Brain cortex tissue (50 mg) (*n*=3 per genotype) was used to isolate nuclei with the Singulator S100 system. S2 Genomics cartridges (S2 Genomics, #100-063-287) were pre-chilled to-20°C before the start of the protocol. Conical tubes (15 mL) and the singulator equipment were pre-chilled before the start of the isolation. The right cortex tissue was placed inside the pre-chilled cartridge with 75μL of RNase inhibitor (Millipore Sigma, 3335399001). Samples were processed as previously described [89]. The resulting nuclear suspension was then transferred to 15 mL tubes and centrifuged at 500xg for 5 minutes at 4°C in a spinning-bucket rotor centrifuge to pellet the nuclei. The supernatant was aspirated and the pellet was subsequently resuspended in 3 mL of 20% nuclei Debris Removal Stock solution (S2 Genomics, #100-253-628) in Nuclear Storage Reagent (S2 Genomics, #100-063-623). This nuclear suspension was centrifuged 8 minutes at 700xg at 4°C with a brake setting of 3 (3 out of 10, Sorvall Legend X1R Centrifuge (ThermoFisher)). After centrifugation, the gradient was carefully aspirated with a P1000 pipette to remove myelin debris (visible as haze/flakes on top of the gradient). Finally, nuclei were resuspended in 3 mL of resuspension buffer (2% BSA in Nuclear Storage Reagent).

### Library preparation and Sequencing

Nuclei concentration and integrity were assessed via trypan blue staining. Nuclear suspensions were processed with 10x Chromium. Each nuclear suspension was counted and diluted to 1,000 nuclei per μL, then subjected to library preparation on Chromium Next GEM Single Cell 3′ v4 GEM-X dual index kit (10x Genomics) according to the manufacturer’s protocol. The cDNA and library quality were assessed via a 2100 Bioanalyzer (Agilent Technologies). Sequencing was performed on a NovaSeq 6000 (v1.5 S2; Illumina).

### snRNA-seq data analysis

The data were processed with the Cell Ranger pipeline (v9.0.1, 10x Genomics) and aligned to GRCm38 [90]. The barcodes, features and count matrix generated by cellranger were loaded into RStudio (v2023.09.1) running R (v4.3.1). The transgene sequences for the Danish amyloid and the *Mapt^-/-^* replacement for Neomycin Resistance (pNeoR) were created as custom references and added to the GRCm38 data. SoupX (v1.6.2) was used to quantify ambient RNA and clean the data [91]. Doublets were identified and removed with DoubletFinder (v2.0.6) [92]. The data was then loaded into Seurat (v5.0.2) [93]. Data was normalized using SCTransform (v0.4.1) [94]. Nuclei were then clustered via the first 20 principal, selected based on the elbow plot. Clusters were annotated using Enrichr [95]. Differences in cell type proportions were quantified using speckle (v1.2.0) [96]. Inference of cell-cell communication from interacting cells groups was performed with CellChat v2 (v2.1.0) [97].

### Human iPSC Bulk RNA-seq analysis and overlap analysis

Bulk RNA-sequencing data from human iPSC-derived glutamatergic neuronal cultures treated with tau antisense oligonucleotides (Tau ASO) or non-targeting ASO controls were obtained from NCBI Gene Expression Omnibus (GEO accession number GSE204931). Sequencing data was processed using the previously reported alignment-based workflow [98]. Briefly, reads were aligned to the human reference genome (GRCh38) using STAR. Gene-level quantification was performed with RSEM using Ensembl gene annotations. Differential gene expression analysis between Tau ASO and control conditions was conducted in R using DESeq2 [99]. Differentially expressed genes were defined using a fold-change threshold of |log2(fold change)| ≥ 0.58 (corresponding to a 1.5-fold change) and a Benjamini-Hochberg (BH) adjusted p-value < 0.05. To enable cross-species comparison with mouse snRNA-seq data, human differentially expressed genes were mapped to their mouse orthologs using g:Profiler [100]. The resulting mouse ortholog gene sets were used to assess concordance with cell-type transcriptional signatures identified by snRNA-seq. Cell-type transcriptional signatures were defined using the Wilcoxon rank-sum test as implemented in Seurat’s *FindMarkers* function. Genes were tested using a minimum expression threshold of 10 cells per group. Differentially expressed genes (DEGs) were defined using a BH-adjusted p-value < 0.05, without applying a log-fold change threshold during testing. Directional DEG sets were subsequently defined based on the sign of the average log2 fold change, with positively enriched genes representing transcripts upregulated in the target cell class relative to all other cells. For each cell class, snRNA-seq derived DEG sets were compared against Tau ASO and non-targeting ASO gene sets using Fisher’s exact test (2 × 2 contingency tables defined by gene membership in the snRNA-seq DEG set and the corresponding ASO RNA-seq DEG set) [101]. The background gene universe was defined as all genes detected in the snRNA-seq dataset (non-zero expression in at least one nucleus), thereby controlling for gene detectability and minimizing bias arising from platform-specific expression constraints. Fisher’s exact test yielded odds ratios and p-values (corrected for multiple comparisons via BH), with enrichment interpreted as a statistically significant overrepresentation of shared genes relative to random expectation. This analysis enabled systematic evaluation of the extent to which cell-type specific transcriptional programs observed in mouse brain tissue recapitulate tau-dependent transcriptional changes in human neuronal cultures.

### Cosine similarity analysis of astrocyte subclusters to previously reported gene programs

To quantify the similarity between published astrocyte gene programs by *Saddick et al* [102] and astrocyte subclusters identified in our dataset, we first identified marker genes for each subcluster using Seurat’s *FindAllMarkers* with the following parameters: logfc.threshold = 0.25, min.pct = 0.1, and only.pos = TRUE. For each subcluster, average expression values of these marker genes were calculated across all cells in the cluster, and cosine similarity was computed between the resulting expression vector and an idealized presence vector (all ones) using the proxy package (v0.4-27). To assess statistical significance, we generated a null distribution by repeating the similarity calculation with the same number of randomly selected genes (excluding the compared genes in program) over 1,000 permutations. Empirical p-values were computed as the proportion of null similarities equal to or exceeding the observed value and were corrected for multiple testing using the Benjamini-Hochberg procedure. This analysis was repeated for each unique gene set within the *Saddick et al*. astrocyte programs. Similarity scores were visualized as a dot plot displaying cosine similarity (color scale) and statistical significance (dot size,-log10 adjusted p-value) for each program-subcluster pair.

### Three-way differential expression analysis and radial visualization (3D Volcano)

To identify and visualize genotype-dependent transcriptional changes within specific brain cell types, we implemented a three-way differential expression framework adapted from the *volcano3D* R package (v2.0.9) [103]. Briefly, the Seurat object was subset by cell type × genotype combinations (e.g., Astrocytes-WT, Astrocytes-Tg-FDD, Astrocytes-Tg-FDD;*Mapt*, and scaled expression values were extracted for the cell type of interest. Three pairwise contrasts were computed between genotypes (A vs. B, B vs. C, and C vs. A).

For each contrast, Seurat’s *FindMarkers* function was used to calculate pairwise differential expression without thresholds on minimum percent expression or log fold change (min.pct = 0, logfc.threshold = 0), returning p-values, adjusted p-values (Benjamini-Hochberg), and log₂ fold changes for all expressed genes. In parallel, an analysis of variance (ANOVA) was applied to each gene across the three genotype groups to test for overall expression differences, while including individual sample identifiers as a covariate to reduce pseudoreplication. Overall p-values were corrected for multiple testing using the qvalue method. Pairwise and overall statistics were integrated into a unified results table and used to assign each gene to one of seven mutually exclusive significance patterns, reflecting genotype-specific or shared effects across conditions. Importantly, gene classification was determined by the combined pattern of pairwise significance and directionality across all three contrasts, rather than by any single adjusted p-value alone. These categories included non-significant (“ns”) genes and genes selectively altered in one genotype or shared between two genotypes [e.g., “Tg-FDD;*Mapt*^-/-^” “Red” (R), “Tg-FDD;*Mapt*^-/-^+WT” “Yellow” (Y), “WT” “Green” (G), “Tg-FDD+WT” “Cyan” (C), “Tg-FDD” “Deep Blue” (DB), “Tg-FDD+Tg-FDD;*Mapt*^-/-^” “Purple” (P)]. Classification and visualization were performed using volcano3D*’s polar_coords* function, which maps genes into polar coordinate space to capture both the magnitude and specificity of genotype effects.

## RESULTS

### snRNA sequencing reveals changes in excitatory neurons in aged tau knockout mice

To investigate the cellular and molecular consequences of tau ablation in a physiological context, we performed single-nucleus RNA sequencing (snRNA-seq) on cortical tissue from 22-month-old wild-type (WT) and tau knockout (*Mapt^⁻/⁻^*) mice (*n*=3). At this age, mice meet the criteria for an elderly phenotype, characterized by widespread senescent alterations across multiple biomarkers [104]. Nuclei were isolated via Singulator S100 and processed via 10x Chromium. After quality control, doublet removal, batch effect correction, and SCT normalization, we obtained 76,036 single-cell transcriptomes. Integrated analysis of all nuclei revealed a well-resolved clustering of major neuronal and non-neuronal cell types (**Fig. 1A**). Separate UMAP projections for WT and *Mapt^⁻/⁻^* samples highlighted subtle genotype-specific shifts in cluster distribution (**Fig. 1A**). Cell-type annotations were validated by the expression of canonical markers for excitatory and inhibitory neurons, astrocytes, oligodendrocytes, immune cells, and vascular-associated cells (**Supplementary Fig. 1A**). Given that most of the literature on tau ablation in pathological and non-pathological contexts describe alterations in neuronal populations [10, 22, 23, 41, 105], we first focused our analysis on neurons to determine whether similar changes occurred in our model by performing a cell type proportion analysis between WT and *Mapt^⁻/⁻^*mice in both excitatory and inhibitory neurons (**Fig. 1B-C**). Accordingly, we observed alterations in a specific cluster of excitatory neurons in *Mapt^⁻/⁻^* mice (**Fig 1B**) while no significant changes in inhibitory neurons proportions were observed (**Fig 1C**). The proportion of cells in Excitatory Neuron 12 (ExNeuron12) was uniquely expressed in the *Mapt^⁻/⁻^* model compared to wild type. To investigate whether these changes in abundance were associated with alterations in cellular signaling, we performed CellChat analysis specifically on the ExNeuron12 cluster. This revealed distinct differences in glutamatergic signaling, with impact in both outgoing and incoming interactions involving ExNeuron12 (**Fig. 1D**). Outgoing interactions represent ligand expression by ExNeuron12 neurons acting on receptors in neighboring cell populations, whereas incoming interactions reflect receptor engagement in ExNeuron12 neurons driven by ligands expressed by other cells. Interestingly, ExNeuron12 exhibited a pronounced enrichment in outgoing glutamatergic signaling, characterized by glutamate ligand activity driven by *Slc1a2* + *Gls* acting on the AMPA receptor subunit *Gria3*, alongside a strong incoming interaction involving glutamate signaling onto the metabotropic glutamate receptor *Grm1* (**Fig. 1D**). In this framework, outgoing interactions reflect glutamate release and signaling from ExNeuron12 neurons to surrounding cells, whereas incoming interactions capture glutamatergic inputs received by ExNeuron12 from the local cellular environment. To contextualize these findings, **Figure 1E** illustrates the established role of glutamate transporters in regulating synaptic glutamate homeostasis. Under physiological conditions, SLC1A2 mediates the majority of glutamate clearance from the synaptic cleft, thereby limiting excessive glutamate activation, preventing excitotoxicity. In contrast, reduced or impaired SLC1A2 function leads to glutamate accumulation in the synaptic cleft, sustained receptor activation, calcium influx, and excitotoxic neuronal damage [106, 107]. Together, these findings suggest that the altered outgoing and incoming glutamatergic communication observed for ExNeuron12 is consistent with synaptic homeostasis and protection against excitotoxic stress.

**Fig 1:**
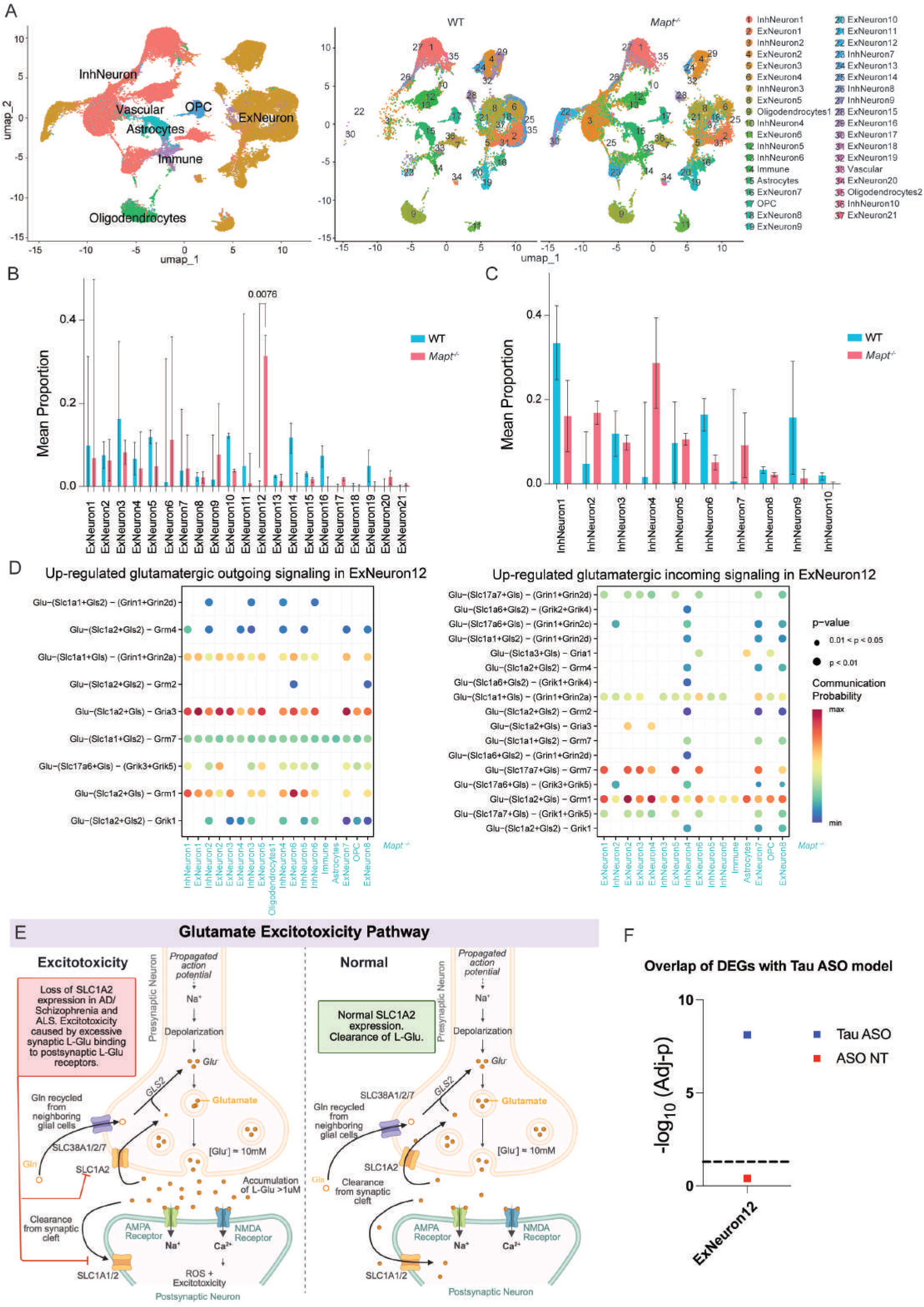
Ablation of endogenous *Mapt* reshapes excitatory neuron composition and recapitulates tau-dependent molecular signatures observed in Tau ASO human neurons. **A.** Uniform Manifold Approximation and Projection (UMAP) of snRNA-seq analysis of 76,036 cells from WT and *Mapt^-/-^* mouse cortex (left panel) and across genotypes (right panel) (*n*=3 males per genotype). **B.** Barplot graphs depicting cell-type proportion analysis comparing WT and *Mapt^-/-^* excitatory neurons. **C.** Barplot graphs depicting cell-type proportion analysis comparing WT and *Mapt^-/-^* inhibitory neurons. Data is shown as mean + SEM, t-test after logit-transformation. **D.** Top outgoing (left panel) and incoming (right panel) upregulated ligand-receptor communications in the glutamate signaling pathway in ExNeuron12. **E.** Schematic illustrating the role of glutamate synthesis, release, receptor activation, and clearance in excitatory neurotransmission. Under AD, schizophrenia, and ALS reduced or impaired SLC1A2 function leads to glutamate accumulation in the synaptic cleft, sustained activation of ionotropic and metabotropic glutamate receptors, increased intracellular calcium, and excitotoxic neuronal damage, leading to excitotoxicity. Under physiological conditions, SLC1A2 mediates the majority of glutamate uptake from the synaptic cleft, limiting excessive receptor activation and preventing excitotoxicity. **F.** Overlap analysis between mouse ExNeuron12 cluster-type specific gene signature and differentially expressed genes from human iPSC-derived glutamatergic neuronal cultures treated with tau antisense oligonucleotides (Tau ASO) or non-targeting ASO controls. Human differentially expressed genes were mapped to mouse orthologs and compared using Fisher’s exact test followed by Benjamini-Hochberg (BH) correction.

To further examine whether molecular changes observed in ExNeuron12 recapitulate transcriptional changes associated with tau modulation in a human model, we integrated transcriptomic data from human iPSC-derived glutamatergic neurons treated with tau antisense oligonucleotides (Tau ASO) or non-targeting ASO controls [98] with our mouse snRNA-seq cell-type signatures (**Fig. 1F**). Human DEGs were mapped to mouse orthologs and compared against cell-type specific gene sets derived from snRNA-seq using Fisher’s exact test. ExNeuron12 displayed a higher overlap with Tau ASO responsive genes (odds ratio = 2.23, Adj-P = 7.62E-09) compared to the non-targeting ASO condition (odds ratio = 1.7, Adj-P =0.39), indicating similarity between a tau reduction strategy in human glutamatergic neurons and the ExNeuron12 transcriptional program identified *in vivo*.

Together, these results identify ExNeuron12 as a selectively altered excitatory neuronal population in the context of tau ablation, characterized by changes in abundance, glutamatergic communication, and transcriptional programs that align with tau-dependent molecular signatures observed in human neurons after tau modulation.

### Endogenous tau ablation promotes astrocytic protective states and glial support of neuronal connectivity

Tau is known to also been express in non-neuronal cells [108–111] (**Supplementary Figure 2A-B**). Astrocytes, as key regulators of synaptic homeostasis, metabolic support, and neuroinflammation, are particularly sensitive to changes in the neuronal environment and have been implicated in numerous neurodegenerative disease processes [112–114]. To elucidate the potential astrocytic alterations in the absence of tau we performed differential gene expression analysis on the entire astrocyte population (4,162 astrocytes), comparing WT and *Mapt^⁻/⁻^* samples. The resulting volcano plot (**Fig. 2A**) highlights a set of genes significantly altered in *Mapt^⁻/⁻^* astrocytes, revealing both upregulated (757, blue DEGs) and downregulated transcripts (248, orange DEGs). Functional enrichment of the upregulated DEGs in astrocytes indicated that tau absence modulates pathways related to synaptic signaling, axon guidance, and ion transport, suggesting altered neuron-astrocyte interactions and potential shifts in astrocytic support of excitatory neurotransmission, an adaptation that may impact gliovascular and overall neuronal function (**Fig. 2B**). Furthermore, we identified 8 subclusters (Subclusters 0-7; **Fig. 2C**) within all astrocytes from WT and *Mapt^⁻/⁻^* mice. This approach allowed us to resolve distinct transcriptional states, assess genotype-specific distributions, and identify molecular pathways altered in *Mapt^⁻/⁻^* astrocytes. To validate the identities of the eight astrocyte subclusters, we examined the expression of canonical astrocytic markers, including *Slc1a2*, *Slc1a3*, *Htra1*, *Luzp2, Gja1, Ctsb, Syt11, Glul, Ptn, Lamp1, S100b, and Cts3* (**Supplementary Fig. 2C**). Astrocyte proportion analysis between WT and *Mapt^⁻/⁻^* revealed that *Mapt^⁻/⁻^*-derived cells are significantly enriched in subcluster 4 (**Fig. 2D**). Next we sought to compare their signature with other astrocytes in context of neurodegeneration previously reported in literature to determine if the identified astrocyte subclusters exhibited similar or distinct transcriptional states [102]. The analysis revealed that most clusters adopted previously reported phenotypes (**Fig. 2E**). Notably, subcluster 4 was enriched for a protective phenotype. Other subclusters were associated with synaptic integrity and glutamate signaling signatures. Interestingly, subcluster 5 did not resemble any previously reported astrocytic transcriptional state, suggesting the presence of a potentially novel phenotype. The protective phenotype in subcluster 4 is of particular interest, given that it was significantly enriched in *Mapt^⁻/⁻^*-derived astrocytes. To further characterize the molecular features underlying this ECM + Protective astrocyte state, we visualized the relative expression of representative genes belonging to the ECM + Protective state. The representation shows relative expression across astrocyte subclusters using a row-scaled (z-score) heatmap (**Supplementary Fig. 3**). This analysis highlights the distinct enrichment signature of genes associated with neuroprotective functions relative to other populations, potentially reflecting the activation of heightened protective mechanisms in response to the absence of physiological tau.

**Fig 2:**
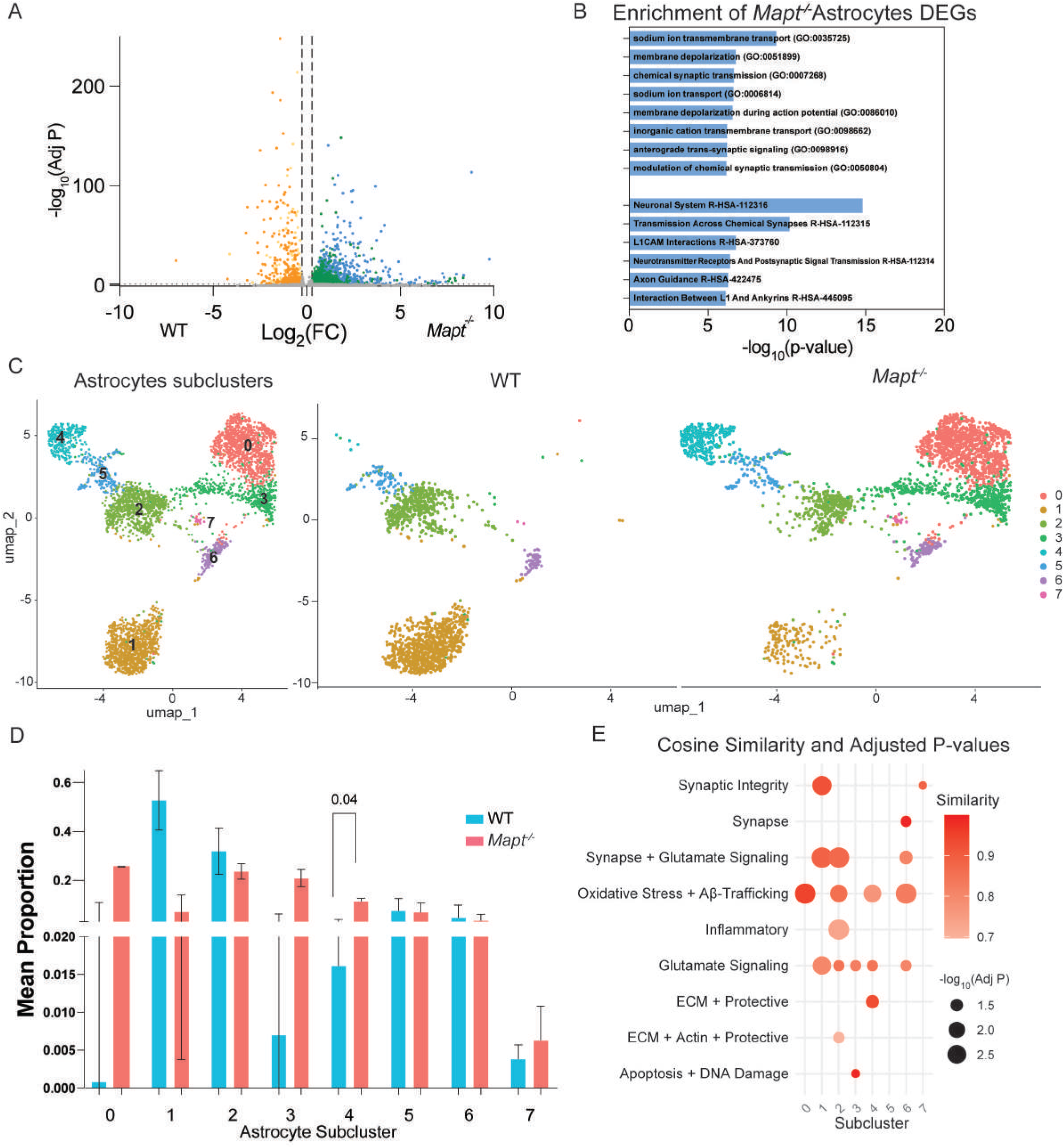
Astrocytes in *Mapt* knockout mice exhibit changes in populations and gene expression profile associated to synapse support and protection **A.** Volcano plot showing DEGs between WT versus *Mapt^-/-^* astrocytes. Discontinued lines parallel to *y-axis* indicates a fold change over 1.5 (log_2_(FC) =±0.58) and discontinued lines parallel to *x-axis* indicates BH adjusted p value of 0.05 (-log_10_ (Adj-P) = 1.3). Upregulated genes are highlighted in Green and Blue if they were also significant after pseudobulking testing; downregulated genes are highlighted in Yellow and Orange if they were also significant after pseudobulking testing. **B.** Enrichment analysis bar graph showing the top biological pathways or processes associated with upregulated *Mapt^-/-^* astrocyte DEGs (Blue DEGs) (Upper bars show GO, lower bars show Reactome). **C.** UMAP representation of astrocyte subcluster (left panel) and across genotypes (right panel). **D.** Barplot graph depicting cell-type proportion analysis comparing WT and *Mapt^-/-^*astrocytes. Data is shown as mean + SEM, t-test after logit-transformation. **E.** Dotplot showing the cosine similarity analysis, which measures how closely two gene expression profiles align, comparing the obtained astrocytic subclusters with previously reported transcriptional signatures of mouse astrocytic subclusters (Sadick, *et al.*, 2022) [102]. The color gradient indicates the degree of similarity, with darker shades representing higher transcriptional similarity between clusters.

Besides astrocytes, oligodendrocytes also play a pivotal role in maintaining brain physiology and responding to pathological conditions [115]. Oligodendrocytes are essential for axonal myelination, metabolic support of neurons, and maintaining white matter integrity [116]. Previous studies have reported expression of tau in oligodendrocytes, both at the mRNA and protein level [117]. Our dataset indicated that tau is expressed highest in oligodendrocytes, supporting the previous findings. Differential gene expression analysis between WT and *Mapt^⁻/⁻^* oligodendrocytes (5,232 cells; **Fig. 3A**), revealed 220 upregulated transcripts (blue DEGs) and 21 downregulated transcripts (orange DEGs). Functional enrichment of the upregulated DEGs indicated that tau absence in oligodendrocytes is associated with pathways involved in neuronal communication and excitability, including neuronal synaptic plasticity, neuron migration, nervous system development, axonogenesis, and chemical synaptic transmission (**Fig. 3B**). Subclustering of oligodendrocytes across WT and *Mapt^⁻/⁻^* mice revealed distinct transcriptional states, as visualized in the UMAP projection (Subclusters 0-4, **Fig. 3C**). Expression of canonical oligodendrocyte markers *Mbp, Mag, Mobp, Mog, Plp1, Sox10, Abca8a, Il33, Dusp11, Opalin, C4b,* and *Ptgds* are in **Supplementary Fig. 2D**. The relative proportions of these subclusters were not found to be significatively altered between genotypes (**Fig. 3D)**, suggesting that there are no genotype-dependent shifts in oligodendrocyte states. Our oligodendrocyte results suggest that this type of glia may respond to tau loss by modulating genes important for neuronal connectivity and function, potentially supporting neuroprotection or compensating for altered neuronal signaling.

**Fig 3:**
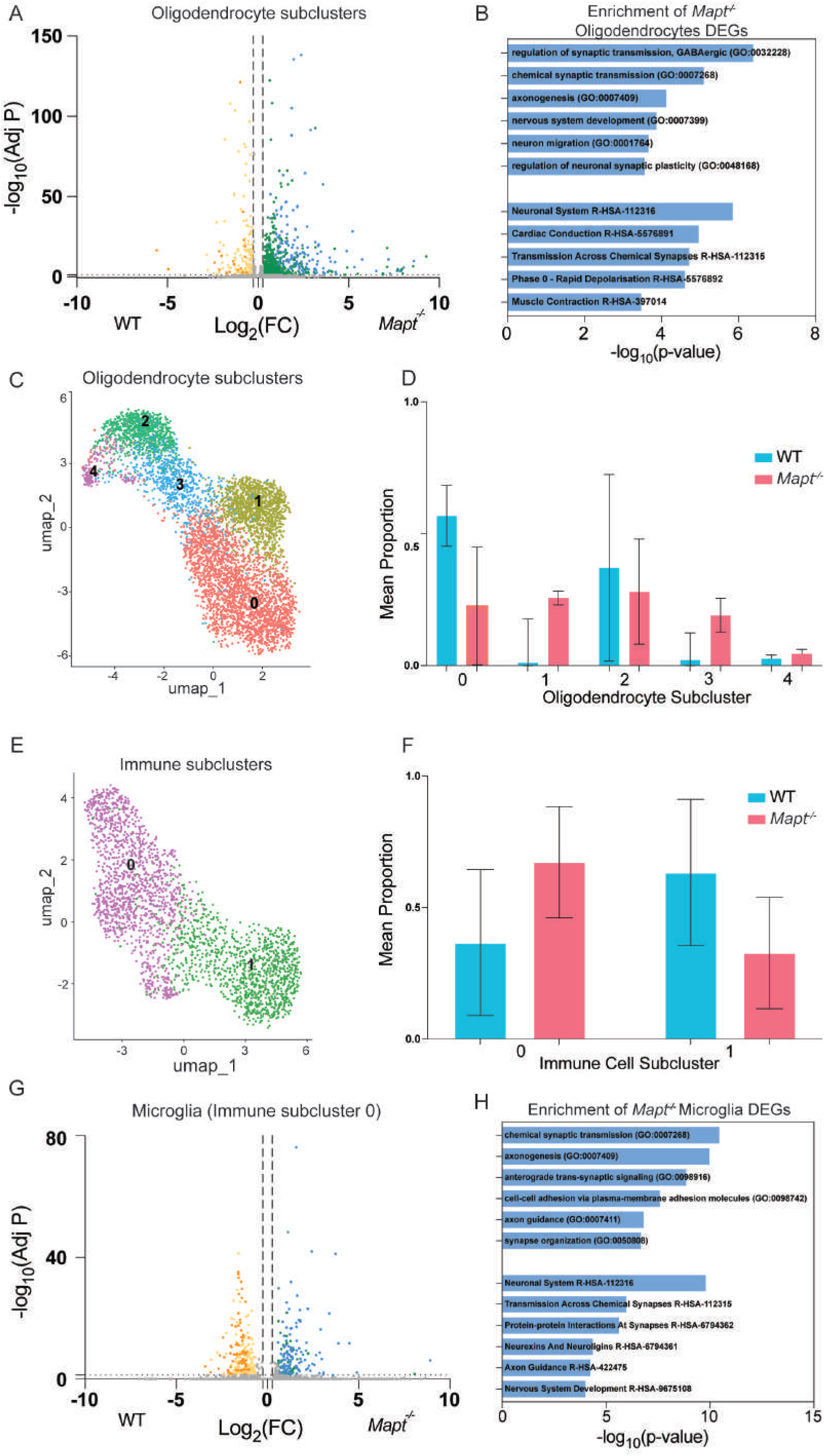
Oligodendrocytes and Microglia cells in *Mapt* knockout mice differentially express genes involved in synaptic function and proteostasis **A.** Volcano plot showing DEGs between WT and *Mapt^-/-^* oligodendrocytes. Discontinued lines parallel to *y-axis* indicate a fold change over 1.5 (log_2_(FC) =±0.58), and discontinued lines parallel to *x-axis* indicates BH adjusted p value of 0.05 (-log_10_ (Adj-P) = 1.3). **B.** Enrichment analysis bar graph showing the top biological pathways or processes associated with upregulated DEGs in *Mapt^-/-^* oligodendrocytes (Blue DEGs). Upper bars show GO, lower bars show Reactome. **C.** UMAP displaying oligodendrocyte subclusters. **D.** Barplot graph depicting cell-type proportion analysis comparing WT and *Mapt^-/-^* oligodendrocytes. Data is shown as mean + SEM, t-test after logit-transformation. **E.** UMAP representing immune cell subclusters. **F.** Barplot graph depicting cell-type proportion analysis comparing WT and *Mapt^-/-^* immune cells. Data is shown as mean + SEM, t-test after logit-transformation. **G.** Volcano plot showing DEGs between WT and *Mapt^-/-^*microglial cells. Discontinued lines parallel to *y-axis* indicates a fold change over 1.5 (log_2_(FC) =±0.58) and discontinued lines parallel to *x-axis* indicates BH adjusted p value of 0.05 (-log_10_ (*p*-value) = 1.3). Upregulated genes are highlighted in Green and if they were also significant after pseudobulking testing, they are highlighted in Blue; downregulated genes are highlighted in Yellow and if they were also significant after pseudobulking testing, are highlighted in Orange. **H.** Enrichment analysis bar graph showing the top biological pathways or processes associated with upregulated DEGs in *Mapt^-/-^* microglia (Blue DEGs) (Upper bars show GO, lower bars show Reactome).

We also examined transcriptional diversity within the immune compartment across WT and *Mapt^⁻/⁻^* genotypes (2,424 cells). We subclustered the immune compartment and analyzed cells that had microglia signatures (Cluster 0; 1,310 cells). Notably, the microglial subpopulation was identified by markers such as *Inpp5d, Hexb, Tgfbr1, and Itgb5* while other immune lineages were identified by expression of *Bcl11a*, *Rora, Psap,* and *Zbtb16* expression (**Supplementary Fig. 2E**). UMAP projection of immune cells allowed visualization of the microglial subcluster (**Fig. 3E**). Comparative analysis of immune cell proportions highlighted that there are no significant genotype-associated shifts, suggesting tau ablation does not impact immune cell composition (**Fig. 3F**). While tau ablation does not impact cell composition, differential gene expression analysis between WT and *Mapt^⁻/⁻^* microglia still revealed significant transcriptional changes, with a set of *Mapt^⁻/⁻^*-specific DEGs (137 Blue Genes) enriched in pathways relevant to synaptic terms, i.e. chemical synaptic transmission, anterograde trans-synaptic signaling, synapse organization, and protein-protein interaction at synapses (**Fig. 3G-H**). These findings indicate that tau loss drives microglia toward a transcriptional state associated with synaptic interaction and structural remodeling, implying modified microglial contributions to synaptic maintenance in the healthy aged brain.

Unlike astrocytes, where a protective transcriptional shift is more directly evident (e.g., increased proportion of a protective astrocyte subcluster), oligodendrocytes and microglia might engage mechanisms that reinforce pathways critical for neuronal support and communication in the absence of tau.

### Impact of tau ablation on intercellular communication

To determine how endogenous tau deletion influences intercellular communication under physiological conditions, we performed ligand-receptor analysis using CellChat, which infers communications based on the coordinated expression of ligands, receptors, and cofactors [97]. Pairwise comparison between WT and *Mapt^⁻/⁻^* showed an overall increase in predicted cell-to-cell communication among most cell types, particularly inhibitory and excitatory neurons in *Mapt^⁻/⁻^*mice (**Fig. 4A**). Cluster-level analysis was then performed using a merged CellChat object containing clusters with inferred interactions in both conditions, enabling a direct comparison of signaling dynamics across shared populations. This analysis highlighted specific cell populations with altered signaling behavior. In particular, InhNeuron2, ExNeuron3, and Oligodendrocytes2 exhibited marked changes in both incoming and outgoing interaction strengths (**Fig. 4B**). For reference, the corresponding cluster-level comparisons without restricting to clusters with inferred interactions in both groups are shown in **Supplementary Fig. 4.**

**Fig 4:**
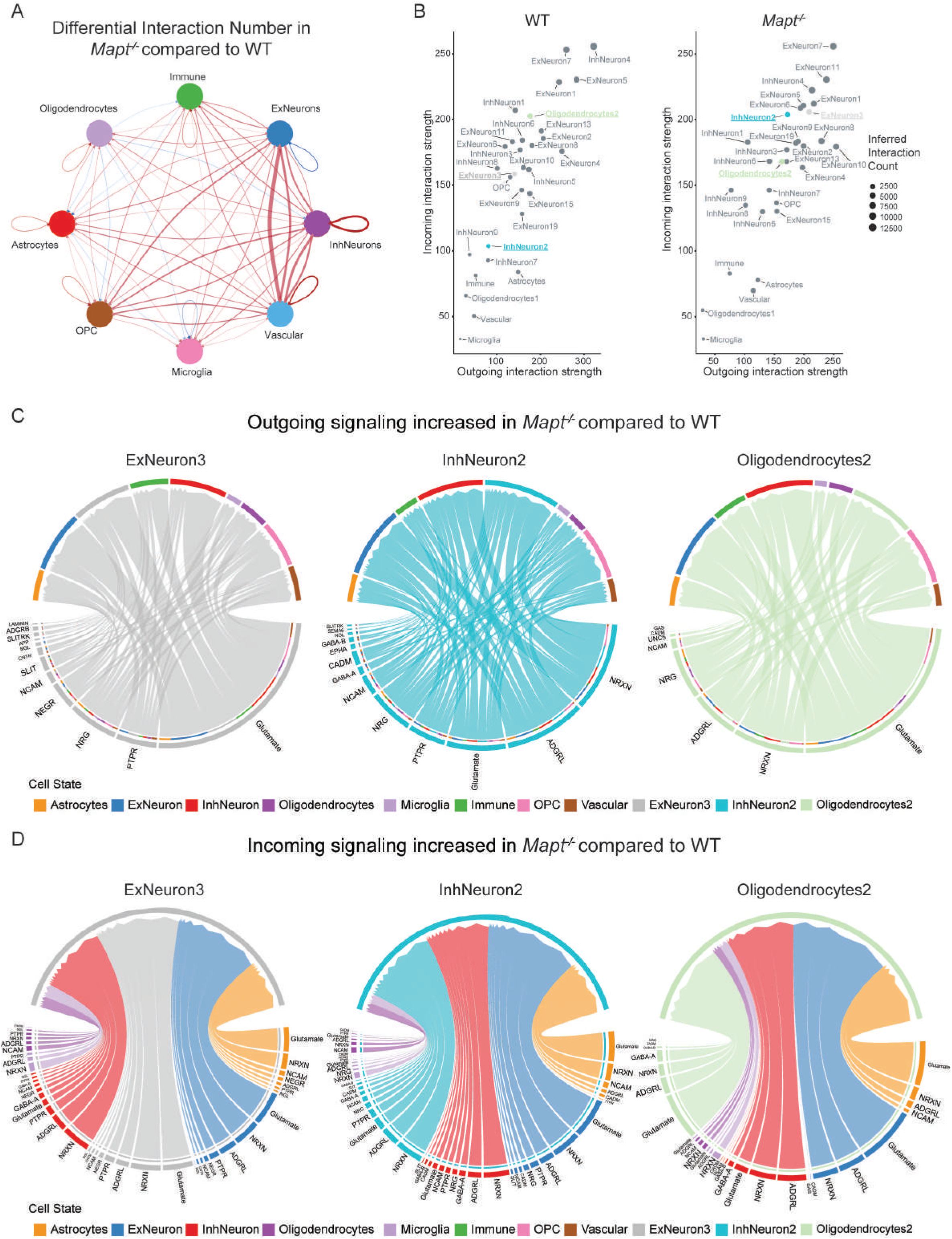
Physiological tau deletion rebalances intercellular communication toward enhanced synaptic signaling. **A.** Chord plot showing the changes in interaction number between the main cell types in the brains of *Mapt^-/-^* mice compared with WT mice. Red lines indicate increased signaling, and blue lines indicate decreased signaling. The thickness of the lines represents the interaction strength. **B.** Predicted interaction strengths of cell clusters with inferred interactions in both conditions, computed from a merged CellChat object for WT (left) and *Mapt^⁻/⁻^*(right). The x-axis shows the interaction strength of outgoing signals, and the y-axis shows the interaction strength of incoming signals. Highlighting specific clusters with altered incoming/outgoing signaling dynamics (ExNeuron3, InhNeuron2, and Oligodendrocytes2). **C.** Outgoing pathway-level signaling chord plots showing increased signaling from ExNeuron3, InhNeuron2, and Oligodendrocytes2 to other cell types in *Mapt^-/-^* compared with WT. **D.** Incoming pathway-level signaling chord plots showing increased signaling to ExNeuron3, InhNeuron2, and Oligodendrocytes2 from other cell types in *Mapt^-/-^* compared with WT.

Pathway-level analysis of these clusters demonstrated increased inferred signaling through synaptic pathways, including Glutamate, NRXN, GABA-A, and GABA-B (**Fig. 4C-D**), based on coordinated ligand-receptor expression patterns [97]. These pathways are central to excitatory and inhibitory neurotransmission, and prior studies have shown that tau reduction modulates neuronal excitability and network synchronization, including suppression of hyperexcitability [22, 76, 81, 118]. Notably, neurexins are presynaptic adhesion molecules critical for synaptic organization and transmission, with additional role in regulating inhibitory neurotransmitter release [119]. Collectively, these findings suggest that physiological tau ablation enhances synaptic and neurotransmitter-mediated intercellular communication.

### Neuronal changes and remodeling of excitatory neuron subtypes after tau ablation in a mouse model of CAA

Having established how tau ablation impacts the aging brain under physiological conditions, we next sought to determine how tau removal influences disease-associated cellular and molecular mechanisms using the Tg-FDD mouse model of CAA, previously characterized by us and others [88, 120]. The Tg-FDD mouse model accumulates Danish amyloid exclusively in the vasculature. Mice present motor impairment, vascular damage, astrocytic reactivity and altered inflammation pathways in a sex dependent manner, with males exhibiting stronger phenotype [85]. We examined genotype-specific transcriptomic landscapes in WT, Tg-FDD, and Tg-FDD;*Mapt^-**/-**^* mouse brains. Separate UMAP visualizations revealed distinct clustering patterns within major cell types for each genotype (total of 10,4157 cells; **Fig. 5A**). Cell type proportion analysis across excitatory and inhibitory neurons showed that only excitatory neurons exhibited significant genotype-dependent changes, with a marked expansion of the Excitatory Neuron 6 (ExNeuron6) cluster in Tg-FDD;*Mapt^⁻/⁻^* brains (**Fig. 5B-C**). Functional enrichment analysis of ExNeuron6 marker genes revealed a strong association with synaptic signaling and plasticity pathway, highlighting a transcriptional program consistent with active synaptic transmission and neuronal connectivity (**Fig. 5D**). To characterize molecular differences in this cluster underlying this expansion, we applied a 3D volcano plot framework to compare high-dimensional gene expression data across genotypes (**Fig. 5E**) [103]. This approach separated DEGs into genotype-specific and shared categories where genes are classified if they are selectively altered in one genotype or shared between two genotypes, identifying a total of five categories: “Tg-FDD;*Mapt*^-/-^” (R), “WT” (G), “Tg-FDD+WT” (C), “Tg-FDD” (DB), “Tg-FDD+Tg-FDD;*Mapt*^-/-^” (P). This analysis identified a set of 20 “Red” genes (R) that were exclusively upregulated in ExNeuron6 from Tg-FDD;*Mapt^⁻/⁻^* mice, absent in both WT and Tg-FDD. To determine if these 20 genes were changed in all excitatory neurons due to genotype, we created a row-scaled (z-score) heatmap of the 20 ExNeuron6-associated (R) genes across genotypes. This revealed selective enrichment of certain genes within Tg-FDD;*Mapt^⁻/⁻^* derived excitatory neurons (**Fig. 5F**). Functional enrichment of these genes indicated that tau absence in ExNeuron6 is associated with pathways related to glutamatergic neurotransmission and postsynaptic signaling, including NMDA receptor activation, regulation of synaptic transmission, anterograde trans-synaptic signaling, and long-term potentiation, suggesting ExNeuron6 may adapt to tau loss by modulating key genes involved glutamate receptor-mediated plasticity (**Fig. 5G**). Together, these findings indicate that tau ablation is accompanied by coordinated transcriptional remodeling in ExNeuron6 that enhances excitatory synaptic signaling and plasticity-related pathways. Notably, these changes do not primarily reflect restoration of disease-associated transcriptional alterations observed in Tg-FDD mice, but instead represent a distinct Tg-FDD;*Mapt^⁻/⁻^*-specific neuronal state characterized by induction of genes not significantly dysregulated in Tg-FDD versus WT conditions.

**Fig 5:**
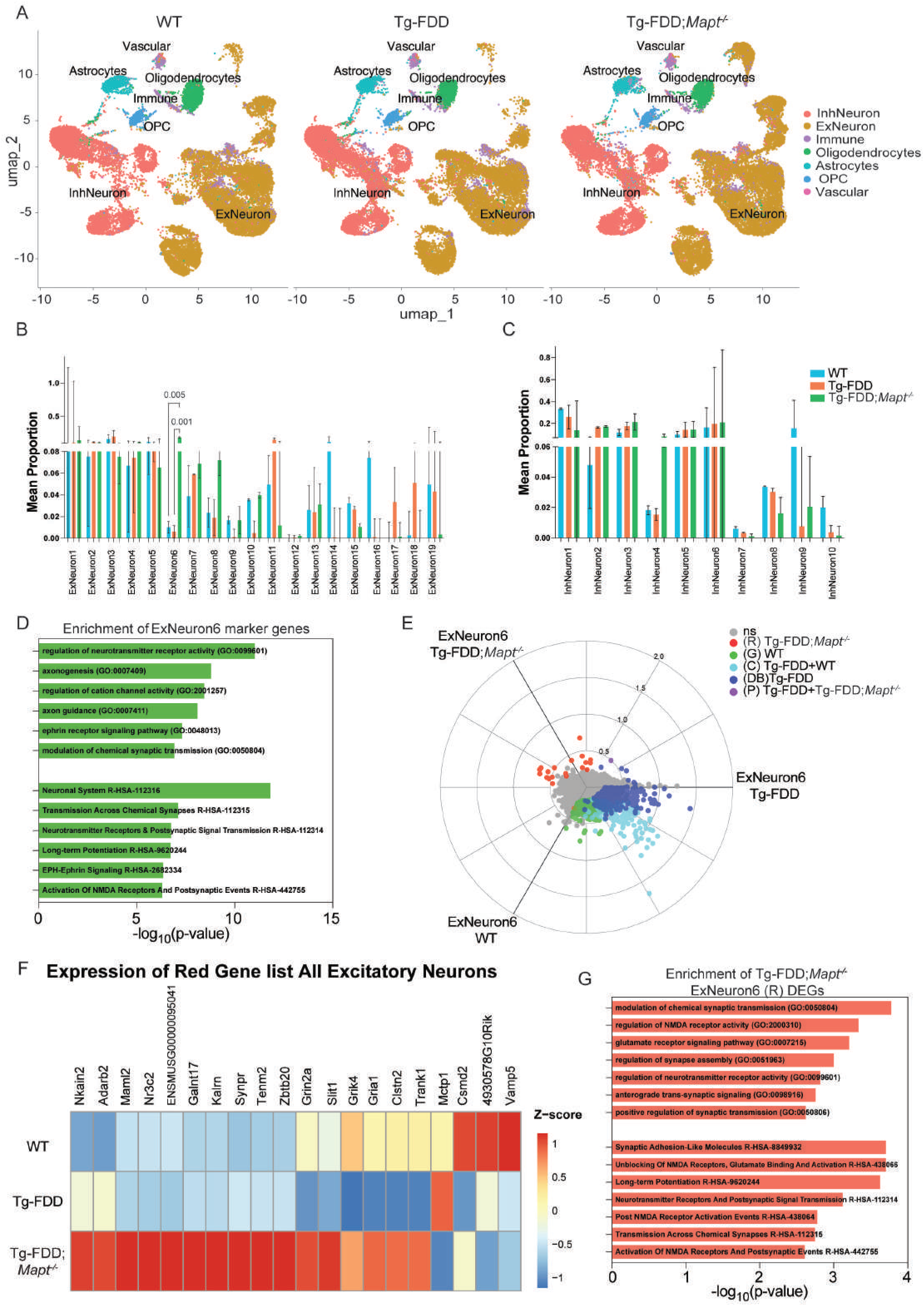
Loss of *Mapt* alters excitatory neuronal populations, synaptic gene expression and intercellular signaling in Tg-FDD model mice **A.** UMAP visualization of snRNA-seq analysis of 104,157 cells from mice cortex from WT, Tg-FDD and Tg-FDD;*Mapt^-/-^*across genotypes. **B.** Barplot graphs depicting cell-type proportion analysis comparing WT, Tg-FDD and Tg-FDD;*Mapt^-/-^* excitatory neurons. **C.** Barplot graphs depicting cell-type proportion analysis comparing WT, Tg-FDD and Tg-FDD;*Mapt^-/-^* inhibitory neurons. Barplots data is shown as mean + SEM, one way ANOVA after logit-transformation, followed by model-based post-hoc pairwise contrasts. **D.** Enrichment analysis bar graph showing the top biological pathways or processes associated with ExNeuron6 cluster markers. Upper bars show GO, lower bars show Reactome. **E.** 3D Volcano plot showing DEGs for ExNeuron6 in a three-way WT, Tg-FDD, and Tg-FDD;*Mapt^-/-^* comparison. Each dot represents a gene. The X-, and Y- axes in the plot show the scaled mean expression differences for each of the three pairwise comparisons between genotypes. Genes further from the origin show larger differential expression. Dot color indicates statistical significance (one way ANOVA followed by BH multiple comparisons tests). Coordinates were calculated using the *polar_coords()* function from the volcano3D package, which enables simultaneous visualization of all three comparisons in one plot. The radical scale axis shows normalized score scale. (R: Red, G: Green, C: Cyan, DB: Deep Blue, P: Purple). **F.** Heatmap of the 20 ExNeuron6-associated (R) genes across genotypes showing the relative expression across all excitatory neurons. Expression values represent average expression per subcluster and are row-scaled (z-scored). **G.** Enrichment analysis bar graph showing the top biological pathways or processes associated with upregulated genes in Tg-FDD; *Mapt^-/-^* ExNeuron6 (R) gene set. Upper bars show GO, lower bars show Reactome.

### Astrocytic transcriptional programs in vascular disease are partially restored by endogenous tau ablation

To determine if astrocytic transcriptional states are altered by endogenous ablation of tau in the context of vascular amyloid pathology, we examined the astrocyte subclusters from the three states (4,104 astrocytes; Subclusters 0-6; **Fig. 6A)**. Here, there was no significant difference in astrocyte subcluster proportions (**Fig. 6B**). Furthermore, global transcriptional changes across the three genotypes using a 3D volcano based differential expression framework were analyzed (**Fig. 6C**). This approach partitioned astrocytic DEGs into genotype-specific and shared categories. As in the ExNeuron6 analysis, the astrocytic analysis identified the “Tg-FDD;*Mapt^⁻/⁻^* unique” (R) gene set, genes whose expression is altered by tau removal but which do not directly contribute to disease-associated transcriptional changes (**Fig. 6D**). In contrast, astrocytic DEGs identified a “Yellow” (Y) gene set that was upregulated in Tg-FDD;*Mapt*^-/-^ and WT but not in Tg-FDD. Notably, genes within this set displayed marked similarity between Tg-FDD;*Mapt*^-/-^, and WT astrocytes, with clear divergence from Tg-FDD astrocytes, highlighting the role of tau removal in restoring specific transcriptional programs toward physiological levels (**Fig. 6E**). Functional enrichment of the Y set revealed pathways involved in neurotransmitter cycling and synaptic support, including glutamate and aspartate transport and uptake. Additional terms pointed to transport processes across the blood-brain barrier and into cells, suggesting preserved vascular-neuronal exchange capacity. These enrichments indicate restoration of metabolic support and extracellular signaling programs that are disrupted in Tg-FDD astrocytes (**Fig. 6F**).

**Fig 6:**
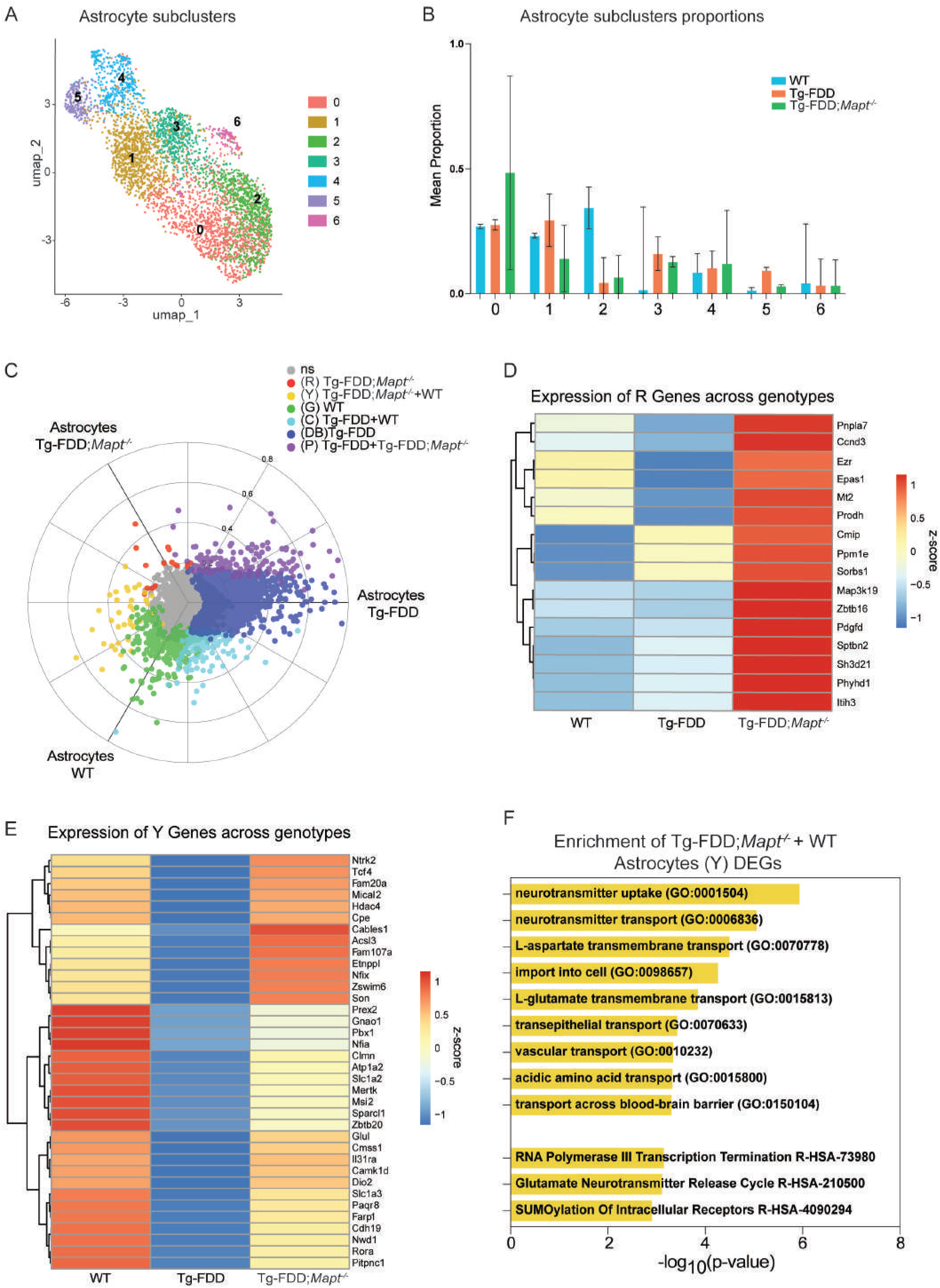
Tau deletion restores astrocytic transcriptional programs in Tg-FDD mice without altering subpopulation composition **A.** UMAP representing astrocyte subclusters. **B.** Barplot graph depicting cell-type proportion analysis comparing WT, Tg-FDD and Tg-FDD;*Mapt^-/-^* astrocytes. Barplot data is shown as mean + SEM, one way ANOVA after logit-transformation**. C.** 3D Volcano plot showing DEGs for astrocytes in a three-way WT, Tg-FDD and Tg-FDD;*Mapt^-/-^* comparison. Each dot represents a gene. The X-, and Y- axes in the plot show the scaled mean expression differences for each of the three pairwise comparisons between genotypes. Genes further from the origin show larger differential expression. Dot color indicates statistical significance (one way ANOVA followed by BH multiple comparisons tests). Coordinates were calculated using the polar_coords() function from the volcano3D package, which enables simultaneous visualization of all three comparisons in one plot. The radical scale axis shows normalized score scale. (R: Red, Y: Yellow, G: Green, C: Cyan, DB: Deep Blue, P: Purple). **D.** Heatmap depicting the Genes that belong to the astrocytic (R) set across genotypes. Row-scaled (z-scored). **E.** Heatmap depicting the Genes that belong to the astrocyte (Y) set across genotypes. Row-scaled (z-scored). **F.** Enrichment analysis bar graph showing the top biological pathways or processes associated with upregulated Tg-FDD;*Mapt^-/-^* & WT astrocytes (Y DEGs: Genes upregulated in both WT and in Tg-FDD; *Mapt^-/-^*, but not in Tg-FDD). Upper bars show GO, lower bars show Reactome.

Together, these results demonstrate that endogenous tau ablation in a vascular disease context selectively restores astrocytic transcriptional programs toward a WT-like state without broadly altering astrocyte subpopulation composition.

### Oligodendrocyte and Microglia cell responses are partially restored to WT levels on endogenous tau ablation in a model of vascular disease

To analyze changes in oligodendrocytes in a disease model when tau is absent, we subclustered the oligodendrocytes (5,650 oligodendrocytes) and obtained 4 subclusters (Subclusters 0-3; **Fig. 7A**). No statistically significant differences in oligodendrocyte subcluster proportions were observed across genotypes (**Supplementary Fig. 5A**). Analysis of the global changes across the genotypes with the 3D volcano framework (**Fig. 7B**) yielded six categories.

**Fig 7:**
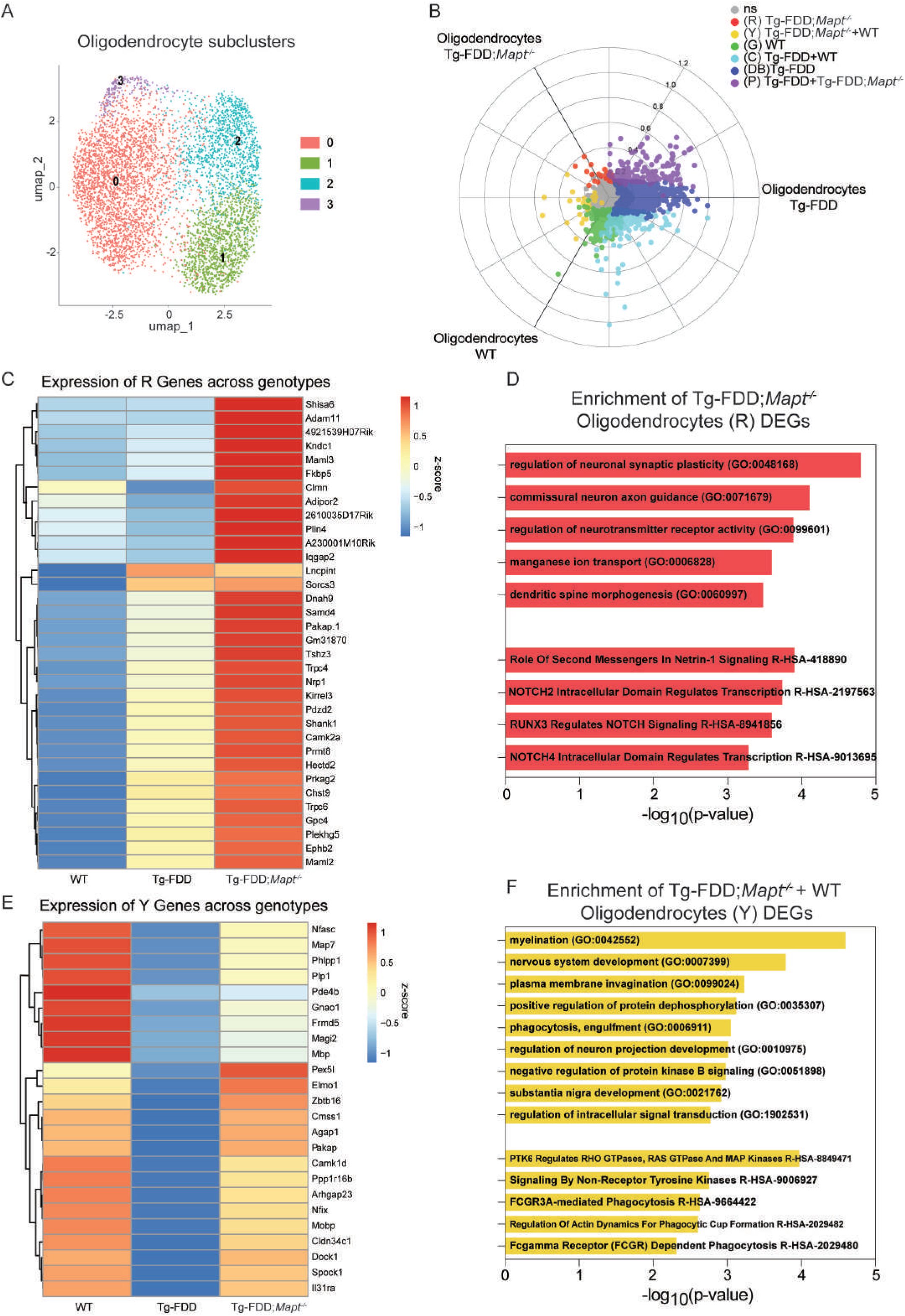
Tau ablation restores oligodendrocyte WT-like transcriptional state in the Tg-FDD mouse model **A.** UMAP representing oligodendrocyte subclusters. **B.** 3D Volcano plot showing DEGs for oligodendrocytes in a three-way WT, Tg-FDD and Tg-FDD;*Mapt^-/-^*comparison. Each dot represents a gene. The X-, and Y- axes in the plot show the scaled mean expression differences for each of the three pairwise comparisons between genotypes. Genes further from the origin show larger differential expression. Dot color indicates statistical significance (one way ANOVA followed by BH multiple comparisons tests). Coordinates were calculated using the polar_coords() function from the volcano3D package. The radical scale axis shows normalized score scale. (R: Red, Y: Yellow, G: Green, C: Cyan, DB: Deep Blue, P: Purple). **C.** Heatmap depicting the Genes that belong to the oligodendrocyte (R) set across genotypes. Row-scaled (z-scored). **D.** Enrichment analysis bar graph showing the top biological pathway s or processes associated with upregulated Tg-FDD;*Mapt^-/-^* oligodendrocytes (R). **E.** Heatmap depicting the Genes that belong to the oligodendrocyte (Y) set across genotypes. Row-scaled (z-scored) **F.** Enrichment analysis bar graph showing the top biological pathways or processes associated with upregulated Tg-FDD;*Mapt^-/-^* & WT Oligodendrocytes (Y) DEGs. Upper bars show GO, lower bars show Reactome.

The (R) gene set across oligodendrocyte genotypes (**Fig. 7C**) illustrates genes selectively altered by endogenous tau ablation that exhibit expression levels exceeding those observed in disease alone. Functional enrichment of the (R) gene set (**Fig. 7D**) revealed pathways related to transcriptional regulation mediated by NOTCH signaling, Netrin-1 signaling, processes associated with dendritic spine morphogenesis, regulation of neuronal synaptic plasticity and neurotransmitter receptor activity. Together, these enrichments suggest reinforcement and support of neuronal connectivity, synaptic modulation, and intracellular transcriptional regulation. Additionally, the (Y) gene set upregulated in both Tg-FDD;*Mapt^⁻/⁻^* and WT but not in Tg-FDD oligodendrocytes (**Fig. 7E**) highlights the role of tau removal in the restoration of specific transcriptional programs toward physiological expression levels. Functional enrichment of the (Y) set (**Fig. 7F**) showed pathways centered on phagocytic activity, including FCGR-dependent engulfment, actin remodeling for phagocytic cup formation, and regulation of RHO/RAS/MAPK signaling. Additional terms encompassed regulation of neuron projection growth, and myelination. All of these suggest that the oligodendrocyte population regains processes for debris clearance, intracellular signaling modulation, and support of neuronal structure and myelin integrity.

The next cellular group that was analyzed was the immune group (4,748 cells). We subclustered the group and obtained 2 subclusters (**Supplementary Fig. 5B**). Cell-type proportion analysis revealed no statistically significant proportion changes across genotypes (**Supplementary Fig. 5C**). Applying the 3D volcano plot framework on the microglia subcluster (subcluster 0; 1383 cells; **Supplementary Fig. 5D**) identified 13 genes associated to (R) set upregulated only in Tg-FDD;*Mapt^⁻/⁻^* and 4 genes in the (Y) set that were selectively upregulated in both Tg-FDD;*Mapt^⁻/⁻^* and WT but not in Tg-FDD microglia. Heatmap visualization of the (R) and (Y) gene sets across genotypes (**Supplementary Fig. 5E-F**) further illustrated shared and distinct transcriptional features between Tg-FDD, Tg-FDD;*Mapt^⁻/⁻^*, and WT microglia, reinforcing the differential impact of tau ablation on disease-independent versus restored transcriptional programs. Together, these findings indicate that changes in oligodendrocytes and microglia with endogenous tau ablation in a vascular disease context are transcriptional rather than compositional, with partial restoration of microglial transcriptional programs toward WT-like states.

### Tau ablation in a disease context reshapes intercellular communications

Having established the effects of endogenous tau deletion under non-disease conditions, we then examined how tau ablation influences intercellular communication in the context of vascular amyloid pathology. We performed the ligand-receptor CellChat signaling analysis comparing Tg-FDD and Tg-FDD;*Mapt^⁻/⁻^* brains. In contrast to the non-disease setting (**Supplementary Fig. 6A**), tau deletion in the disease context resulted in a global reduction in predicted cell-cell communication across most cell types in Tg-FDD; *Mapt^⁻/⁻^* mice (**Fig. 8A**). Cluster-level analysis was used on a merged CellChat object containing clusters with inferred interactions in both conditions.This analysis highlighted InhNeuron4, ExNeuron11, and Astrocytes, which exhibited noticeable changes in both incoming and outgoing interaction strengths relative to Tg-FDD (**Fig. 8B**). Corresponding cluster-level comparisons without restricting to clusters with inferred interactions in both groups are shown in **Supplementary Fig. 6B**. Although the overall inferred communication landscape was reduced, pathway-level chord plots of the previously mentioned clusters revealed persistent selective increases in synaptic and neurotransmitter-related signaling within these clusters (**Fig. 8C-D**). Together, these results suggest that in the presence of vascular amyloid pathology, tau ablation dampens the global intercellular communication network while selectively enhancing synaptic signaling pathways. This context-dependent remodeling of communication networks underscores that the impact of tau deletion differs fundamentally between physiological and pathological environments, likely reflecting compensatory or adaptive mechanisms engaged in response to amyloid-driven network disruption.

**Fig 8:**
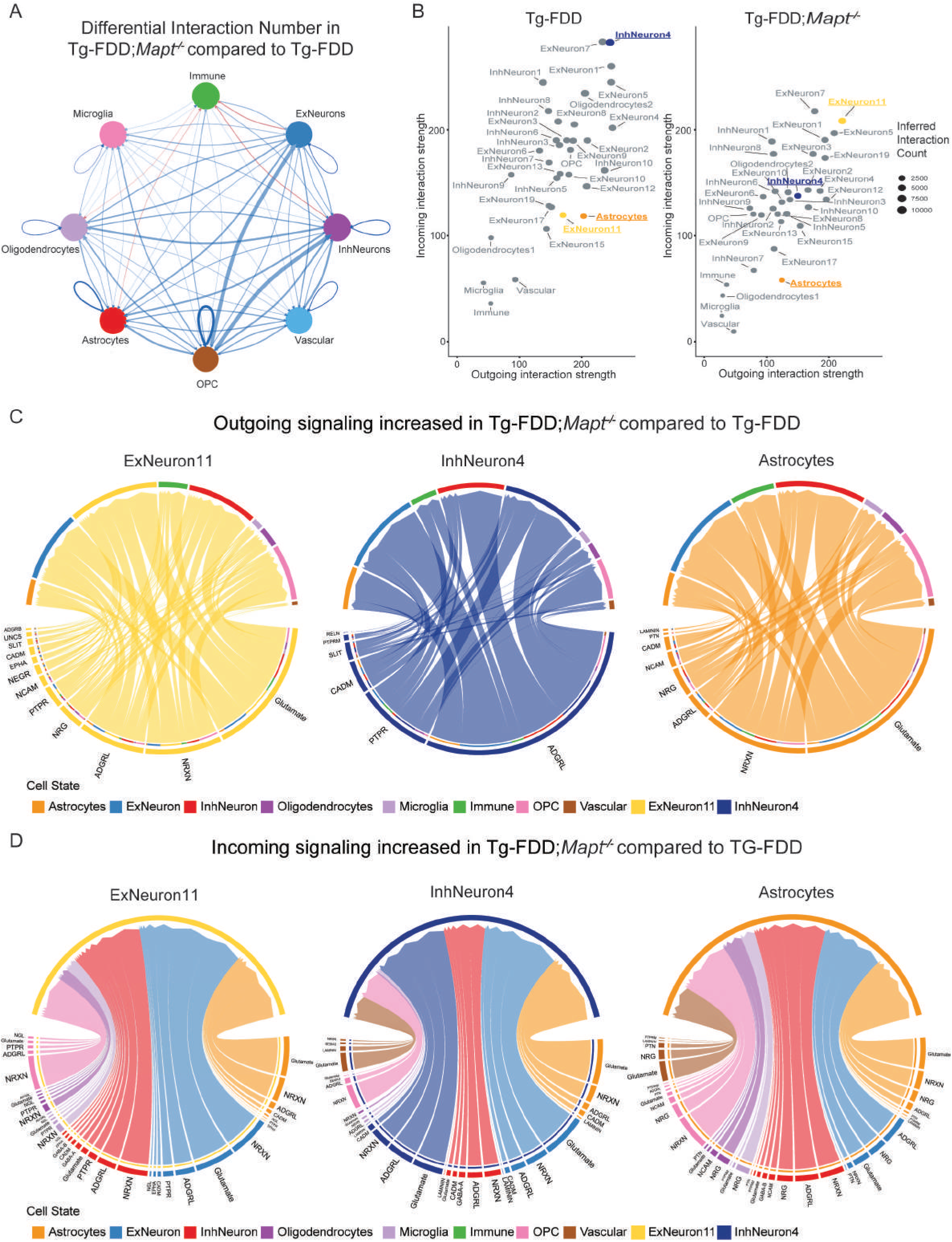
Tau deletion reduces global communication while preserving synaptic signaling in the Tg-FDD mouse model. **A.** Chord plot showing the changes in interaction number between the main cell types in the brains of Tg-FDD;*Mapt^-/-^* mice compared with Tg-FDD mice. Red lines indicate increased signaling, and blue lines indicate decreased signaling. The thickness of the lines represents the interaction strength. **B.** Predicted interaction strengths of cell clusters with inferred interactions in both conditions, computed from a merged CellChat object for Tg-FDD (left) and Tg-FDD;*Mapt^-/-^*(right). The x-axis shows the interaction strength of outgoing signals, and the y-axis shows the interaction strength of incoming signals. Highlighting specific clusters with altered incoming/outgoing signaling dynamics (ExNeuron11, InhNeuron4, and Astrocytes). **C.** Outgoing pathway-level signaling chord plots showing increased signaling from ExNeuron11, InhNeuron4, and Astrocytes to other cell types in Tg-FDD;*Mapt^-/-^*compared with Tg-FDD. **D.** Incoming pathway-level signaling chord plots showing increased signaling to ExNeuron11, InhNeuron4, and Astrocytes from other cell types in Tg-FDD;*Mapt^-/-^*compared with Tg-FDD.

## DISCUSSION

Numerous studies across neurodegenerative and neurodevelopmental models have shown that tau ablation confers neuronal protection resulting in the recovery of behavioral deficits [56, 75, 76, 121–123]. However, it is still unclear mechanistically how tau removal exerts these many protective effects at the cellular and molecular level and which cells are involved in restoring or rewiring transcriptional programs and intercellular communication pathways. Here we identify tau as a context-dependent regulator of brain cellular communication programs, where its ablation induces distinct transcriptional and intercellular rewiring in non-disease aging versus vascular amyloid pathology.

Physiologically, tau knockout (*Mapt^⁻/⁻^*) mice exhibited selective remodeling of excitatory neurons, most prominently an expansion of ExNeuron12. The literature has established a role for tau in regulating glutamatergic synaptic function, where it interacts with Fyn, NMDARs, PSD-95, and SynGAP1 [41, 77, 105]. Through these interactions, tau contributes to the fine-tuning of synaptic transmission and neuronal excitability. However, these same mechanisms may render neurons vulnerable to excitotoxicity, as tau-dependent Fyn-PSD-95-NMDAR coupling amplifies glutamate receptor activation under pathological conditions [52, 124]. In contrast to this excitotoxic amplification, we observed changes pointing to enhanced glutamate clearance capacity through the SLC1A2-GLS axis. SLC1A2 (also known as GLT-1) is the principal mediator of extracellular L-glutamate uptake, predominantly expressed in astrocytes but also present in excitatory neuronal terminals [106, 125]. Loss or dysfunction of SLC1A2 has been implicated in multiple neurodegenerative and neuropsychiatric disorders, including AD where it is closely associated with cognitive decline [126–131]. Our findings suggest a shift from tau-driven excitotoxic amplification toward mechanisms that promote glutamate buffering and synaptic homeostasis, consistent with enhanced SLC1A2-associated glutamatergic regulation. Consistent with this interpretation, conditional SLC1A2 knockout studies have shown that neuronal GLT-1 contributes disproportionately to synaptosomal glutamate uptake relative to its overall protein abundance [132], highlighting a potentially underappreciated role for neuronal SLC1A2 in regulating synaptic glutamate dynamics. Taken together, these observations suggest that removal of endogenous tau not only attenuates pathways that sensitize neurons to excitotoxic stress but also promotes compensatory engagement of glutamate clearance mechanisms, thereby enhancing excitotoxic stress resilience in the absence of physiological tau.

In the context of altered cell abundance and intercellular communication, ExNeuron12 preferentially reflects tau-dependent transcriptional signatures observed in human neurons subjected to targeted tau reduction. The selective enrichment of Tau ASO responsive genes, compared to non-targeting ASO signatures, indicates that this response is specific to tau modulation rather than a generalized effect. This finding is particularly relevant given that current tau-lowering therapies in clinical development broadly target tau expression or accumulation without considering potential cell-type specific adaptations or susceptibilities, even within different neuronal types [133, 134]. By removing tau levels, excitotoxic vulnerability is alleviated. Non-neuronal populations were also affected: astrocytes displayed heterogeneity with the emergence of a subcluster with protective features. This subcluster was enriched for genes involved in extracellular matrix organization [102, 135, 136], metabolic support [135, 137], and inflammatory modulation [138–141]. Genes with participation in processes known to buffer synaptic stress, preserve neuronal integrity, and limit excitotoxic damage were also enriched. These observations align with prior culture studies in which tau reduction shifted astrocytes toward a synaptoprotective state [142]. In the absence of tau, glial populations adopt transcriptional programs consistent with enhanced synaptic support and plasticity. Together, these findings suggest that tau loss in the healthy aged brain engages compensatory and protective mechanisms across multiple cell types.

In contrast, tau removal in a CAA model produced a distinct set of changes. Excitatory neurons responded with an expansion of ExNeuron6. Importantly, evidence suggests that pathogenic gain of function of tau in neurons exacerbates deleterious processes, including overstimulation of PI3K-Akt-mTOR signaling [56]. Tau can regulate this pathway [56], but under pathological stress such as in conditions such as ASD or amyloid accumulation, it can promote network dysfunction [22, 143]. Our data presents the following paradigm; tau ablation alone triggers compensatory remodeling and reduces intrinsic excitotoxic vulnerability. However, in disease settings, this protective effect becomes more evident. Although the amyloid insult persists, removal of tau and its amyloid-induced toxic gain-of-function, together with a rebalancing of excitatory signaling reflected by the enrichment of genes associated with glutamatergic synaptic transmission, enables a shift away from maladaptive processes.

Astrocyte, oligodendrocyte, and microglia subcluster proportions remained stable upon tau loss in the Tg-FDD model, although each exhibited disease-specific transcriptional remodeling. In astrocytes, tau ablation restored a WT-like gene set enriched in neurotransmitter cycling, synaptic support, metabolic exchange, phagocytosis, and signal transduction. These findings suggest a complementary model of benefit: neurons are directly protected by removal of tau’s toxic gain of function, while glial populations likely benefit both from reduced neuronal dysfunction and intrinsic consequences of tau deletion within glial cells. Because tau is eliminated from all cell types in the global knockout, glial transcriptional remodeling may reflect both cell-autonomous and non-cell-autonomous mechanisms whereby relief of neuronal stress reduces maladaptive signaling and enables glia to re-engage adaptive programs [143]. Once tau is eliminated, glia appear better able to mount context and disease-appropriate responses, including elimination of toxic species [144], reduction in kinase activation, and modulation of downstream signaling [76, 78, 145], potentially explaining the broad protective effects of tau ablation across pathological models. Several examples illustrate this targeted restoration. In a vascular injury model induced by high-salt diet, tau ablation rescued molecular pathways underlying neurovascular dysfunction [87]. In the Tg-FDD;*Mapt^⁻/⁻^* model, we previously showed that oligodendrocyte function was restored to WT levels following tau ablation [85]. Notably, recent work has shown that tau neuropathology correlates with hypermethylation of oligodendrocyte genes, leading to reduced expression and demyelination [146]; several of the downregulated genes were recovered in our datasets (*Shank1*, *Clmn*, *Camk1d*, *Mbp*, and *Agap1*), further supporting the link of oligodendrocyte function restoration with endogenous tau removal.

Microglia in the Tg-FDD model showed modest but highly selective transcriptional changes following tau ablation, characterized by small gene sets that distinguished Tg-FDD;*Mapt^⁻/⁻^* microglia from Tg-FDD. This targeted response is consistent with prior reports showing that tau reduction selectively modulates inflammatory and microglial transcriptional programs in amyloid models [147]. Given that microglia can phagocytose extracellular pathological tau [148, 149], and that tau misfolds and acquires toxic gain-of-function properties in our vascular amyloid model [120], tau ablation likely reduces this pathological burden. Pathway analysis highlighted intracellular signaling and metabolic processes, including protein phosphorylation and lipid metabolism, suggesting that tau deletion recalibrates microglial functional states rather than inducing broad inflammatory activation. Notably, components of the TGF-β signaling axis were represented among the regulated genes, consistent with partial restoration of homeostatic regulatory pathways [150–152]. Together, these findings suggest that tau ablation in the vascular amyloid context subtly normalizes microglial signaling competence and homeostatic balance. While neuronal benefits largely reflect cell-autonomous removal of toxic tau functions, glial benefits likely arise from both intrinsic tau loss and relief of external tau-enabled dysregulation.

Finally, intercellular communication further highlighted a clear context-dependent divergence in the impact of tau ablation. Under non-disease conditions, tau loss broadly increased inferred cell-cell communication, particularly among excitatory and inhibitory neuronal populations, consistent with enhanced synaptic and neurotransmission-related signaling. These findings are consistent with a network-level rebalancing and align with emerging evidence that tau modulates global transcriptional programs [153, 154]. In contrast, in the Tg-FDD vascular amyloid context, tau ablation reduced overall predicted intercellular communication while selectively preserving or enhancing synaptic signaling within neuronal and glial clusters. Thus, rather than uniformly amplifying communication, tau deletion appears to remodel signaling networks in a state-dependent manner. It broadly potentiates synaptic interactions in physiology while dampening maladaptive global signaling and refining essential pathways under pathological stress. Taken together, our findings suggest that tau ablation produces two distinct classes of transcriptional responses in the aging brain: disease-independent tau-ablation programs and disease-associated restorative programs. Across multiple cell types, tau removal induced “Tg-FDD;*Mapt^⁻/⁻^* unique” (R) gene sets, representing transcriptional changes that emerged independently of disease-associated dysregulation and were not altered in Tg-FDD versus WT conditions. These programs were particularly prominent in excitatory neuronal populations, where tau ablation promoted novel synaptic signaling and plasticity-associated states rather than simple restoration toward WT expression patterns. In contrast, glia were enriched for (Y) restorative gene sets in which tau ablation re-established WT transcriptional programs disrupted in Tg-FDD mice.

Thus, while neuronal responses to tau loss primarily reflected adaptive remodeling and emergence of new cellular states, glial responses reflected recovery of disease-associated dysfunction.

Importantly, these findings suggest that the beneficial effects of tau reduction are not exclusively dependent on reversing disease pathology, but can also arise through disease-independent compensatory programs induced by tau absence itself. This distinction has important therapeutic implications, as not all neurodegenerative conditions may benefit through identical mechanisms of tau reduction. Instead, the outcome of tau-targeting therapies may depend on the relative contribution of pathology restoration versus tau-ablation-induced adaptive remodeling within specific cell populations and disease environments. More broadly, our study highlights the importance of resolving cell type-specific effects of tau reduction across distinct cell types.

## AUTHOR CONTRIBUTIONS

E.C.-J and C.L-R. designed the study and E.C.-J, N.J.-G., and C.L-R. wrote the manuscript. N.J-G. provided brain samples and performed cortex microdissection. E.C.-J and H.P. performed the snRNAseq experiment. E.C.-J and L.D. performed the data analysis. C.L.-R., E.C.-J, N.J.-G., H.P., L.D. and J.K. critically revised the manuscript and interpreted the data. All the authors have read and approved the final manuscript.

## ACKNOWLEDGMENTS

We would like to acknowledge the Center for Medical Genomics at Indiana University School of Medicine, where the single-nuclei RNA libraries were sequenced, which is partially supported by the Indiana University Grand Challenges Precision Health Initiative and the Indiana Genomic Initiative at Indiana University (INGEN). This research was supported in part by Lilly Endowment, Inc., through its support for the Indiana University Pervasive Technology Institute.

---

**Supplementary Figure 1.**
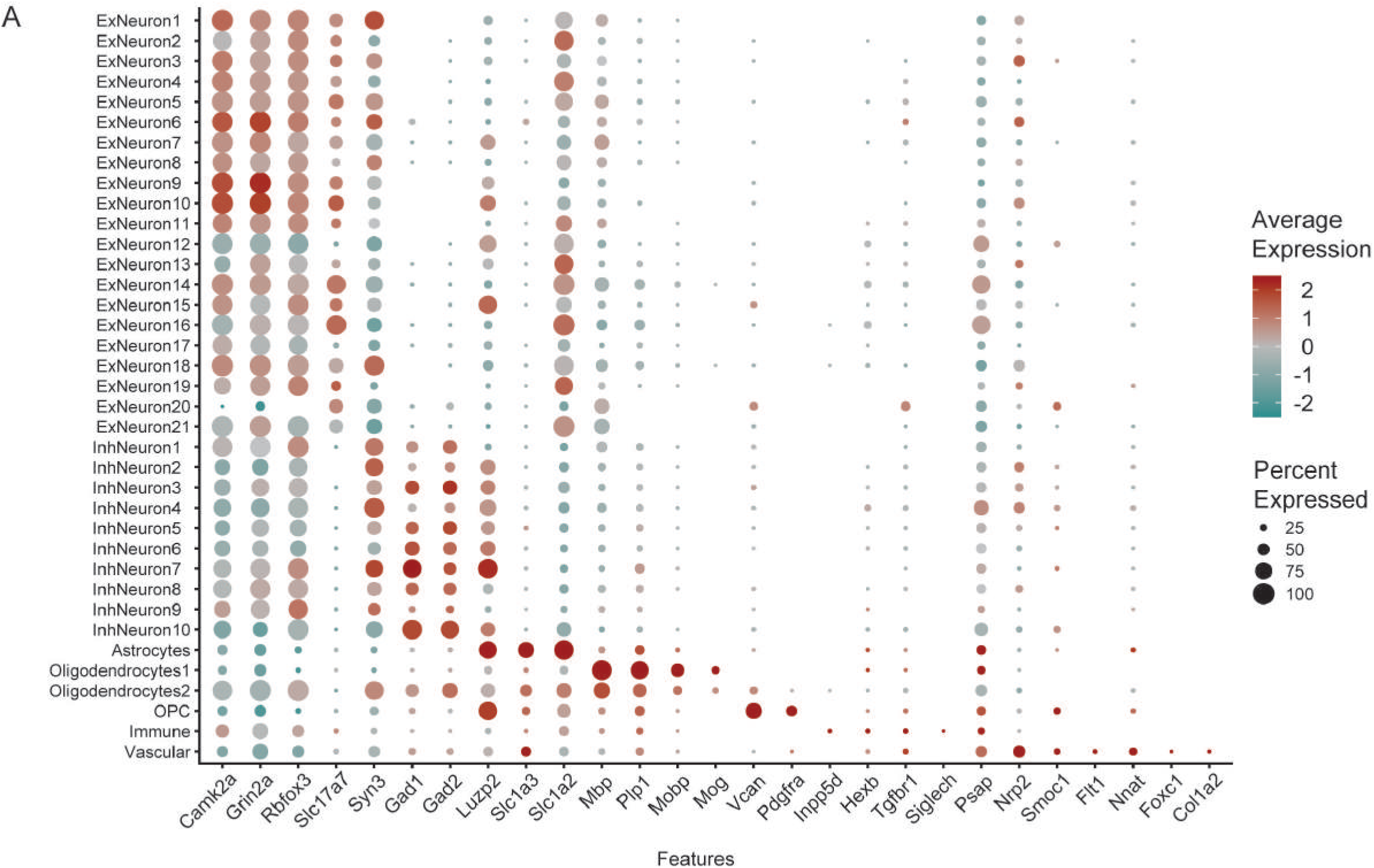
Average scaled expression of canonical markers. **A.** Averaged scaled expression levels of canonical markers genes for different cell types. The size of each dot indicates the proportion of cells within a cluster expressing the gene, the color intensity reflects the average scaled expression level across clusters.

**Supplementary Figure 2.**
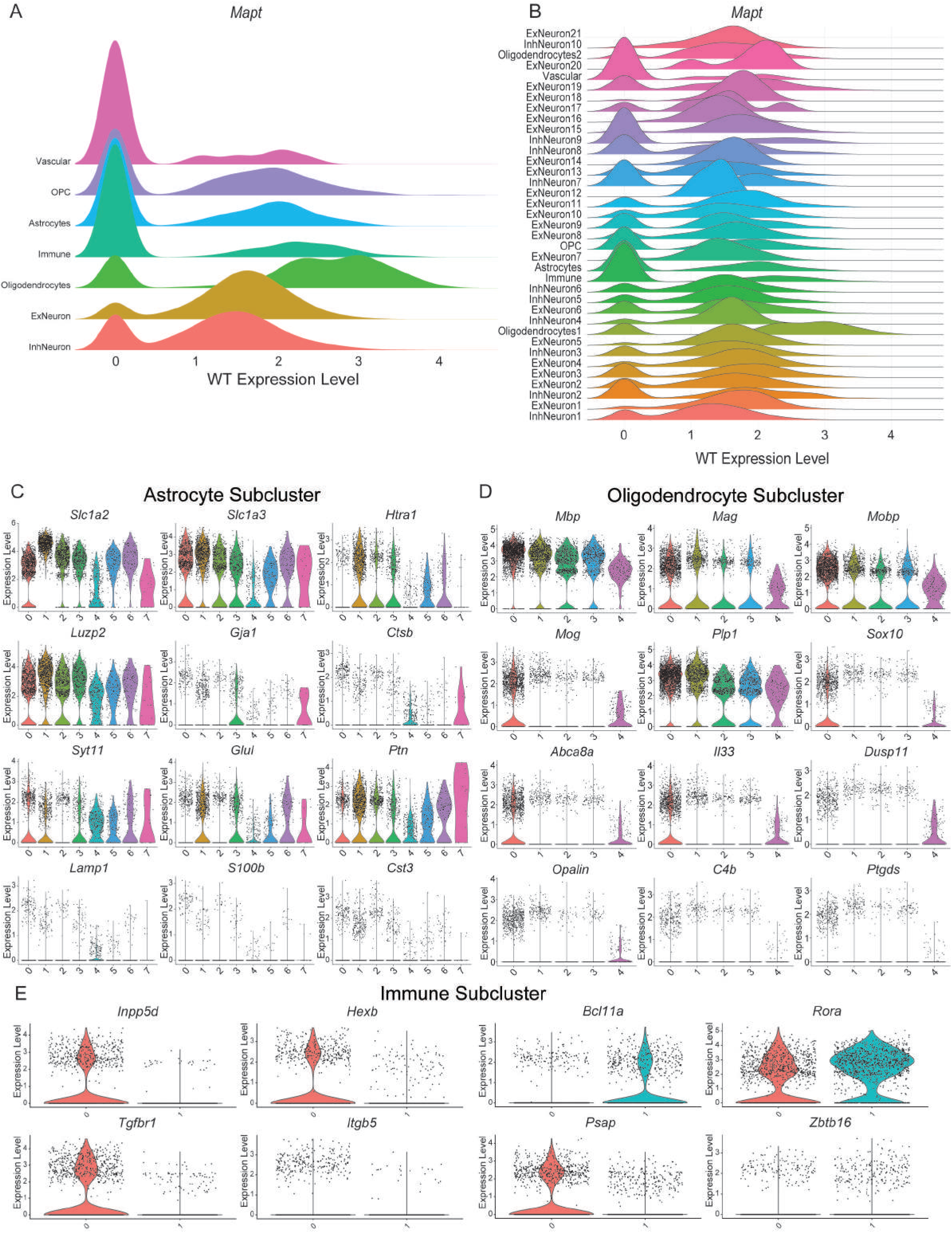
Distribution of *Mapt* expression across cell types and clusters, Subcluster markers **A.** Ridge plot showing the expression levels of *Mapt* across major cell types. Each ridge represents a cell type, and the height of the ridge across the x-axis reflects the number of cells expressing *Mapt* at that level. **B.** Ridge plot showing *Mapt* expression across all detailed cell clusters under the WT condition, highlighting cluster-specific variation in tau levels. **C.** Violin plot showing the expression level of canonical astrocyte marker genes (*Slc1a2*, *Slc1a3*, *Htra1*, *Luzp2, Gja1, Ctsb, Syt11, Glul, Ptn, Lamp1, S100b, and Cts3*) across astrocyte subclusters. **D.** Violin plots showing the expression level of canonical oligodendrocyte marker genes (*Mbp, Mag, Mobp, Mog, Plp1, Sox10, Abca8a, Il33, Dusp11, Opalin, C4b,* and *Ptgds*) across oligodendrocyte subclusters. **E.** Violin plot showing the expression level of marker genes (Microglia: *Inpp5d, Hexb, Tgfbr1, and Itgb5.* Other lineages were identified by expression of *Bcl11a*, *Rora, Psap,* and *Zbtb16*) across immune subclusters.

**Supplementary Figure 3.**
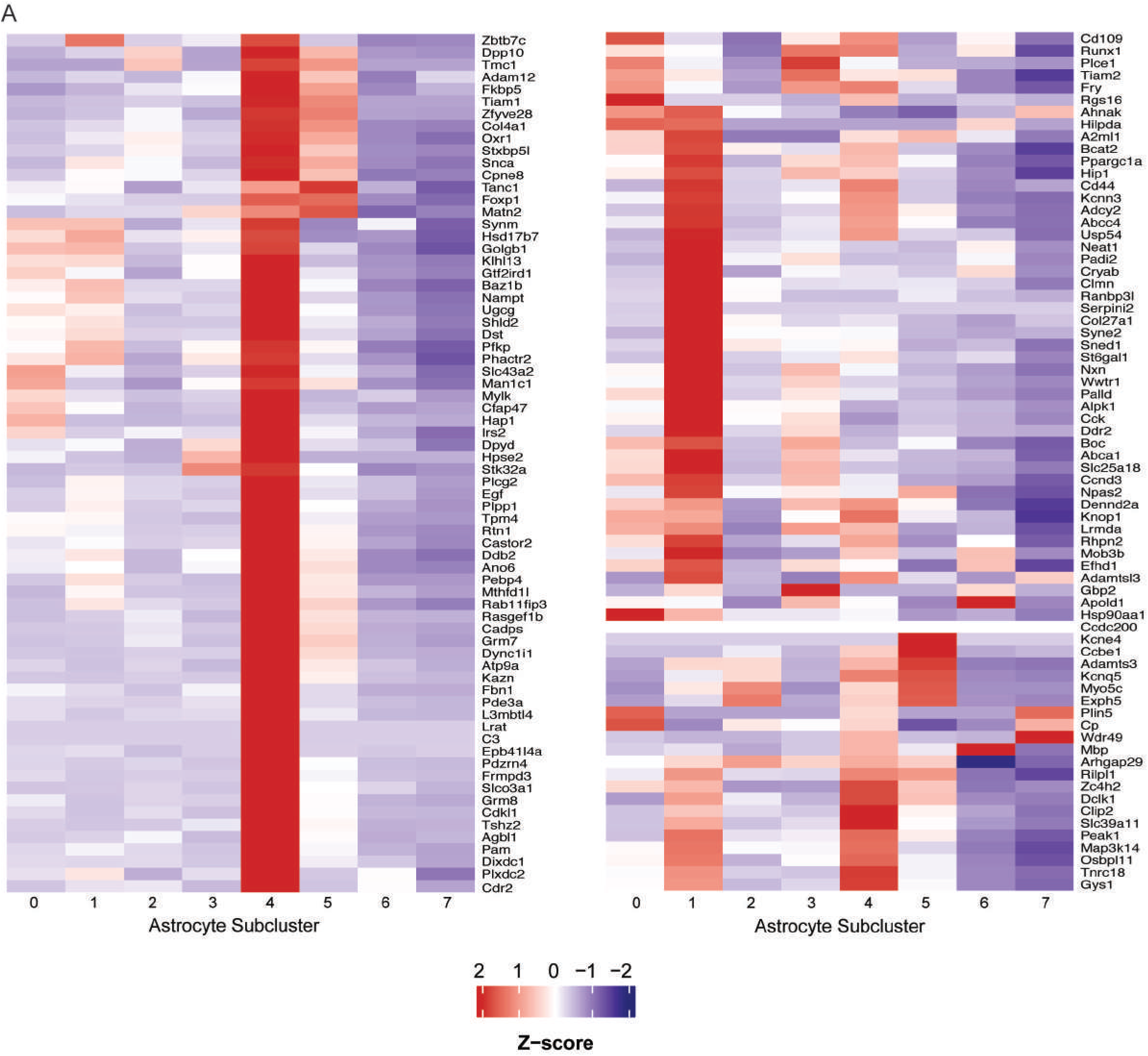
Protective + ECM Heatmap across astrocytic subclusters. **A.** Heatmap showing the relative expression of the entire gene sets markers associated with the ECM and protective signature [102] across astrocyte subclusters. Expression values represent average expression per subcluster and are row-scaled (z-scored).

**Supplementary Figure 4.**
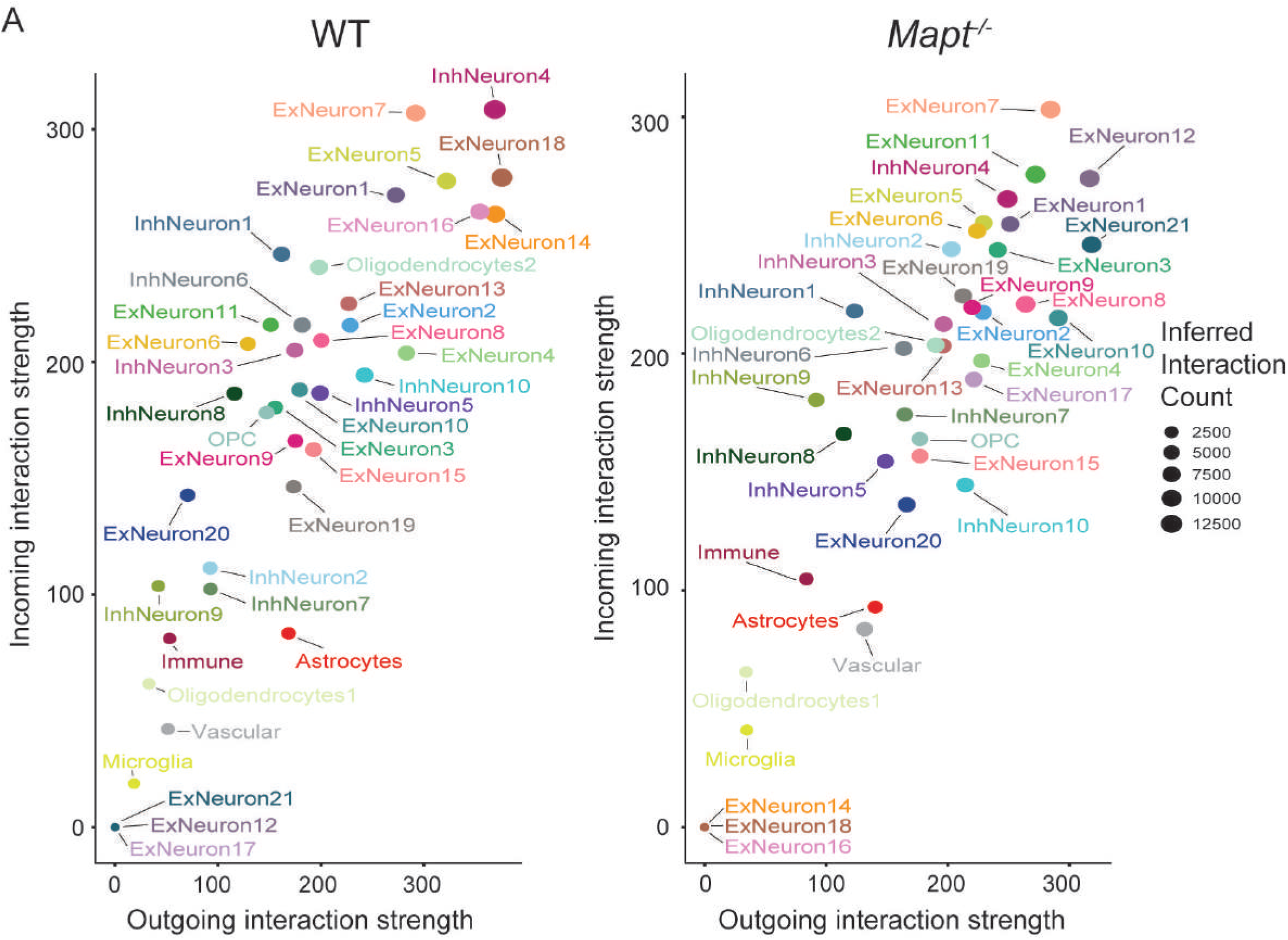
Predicted interaction strengths of physiological tau comparison clusters using CellChat. **A.** Predicted interaction strengths of all cell clusters in both conditions, computed from a merged CellChat object for WT (left) and *Mapt^⁻/⁻^* (right). The x-axis shows the interaction strength of outgoing signals, and the y-axis shows the interaction strength of incoming signals.

**Supplementary Figure 5.**
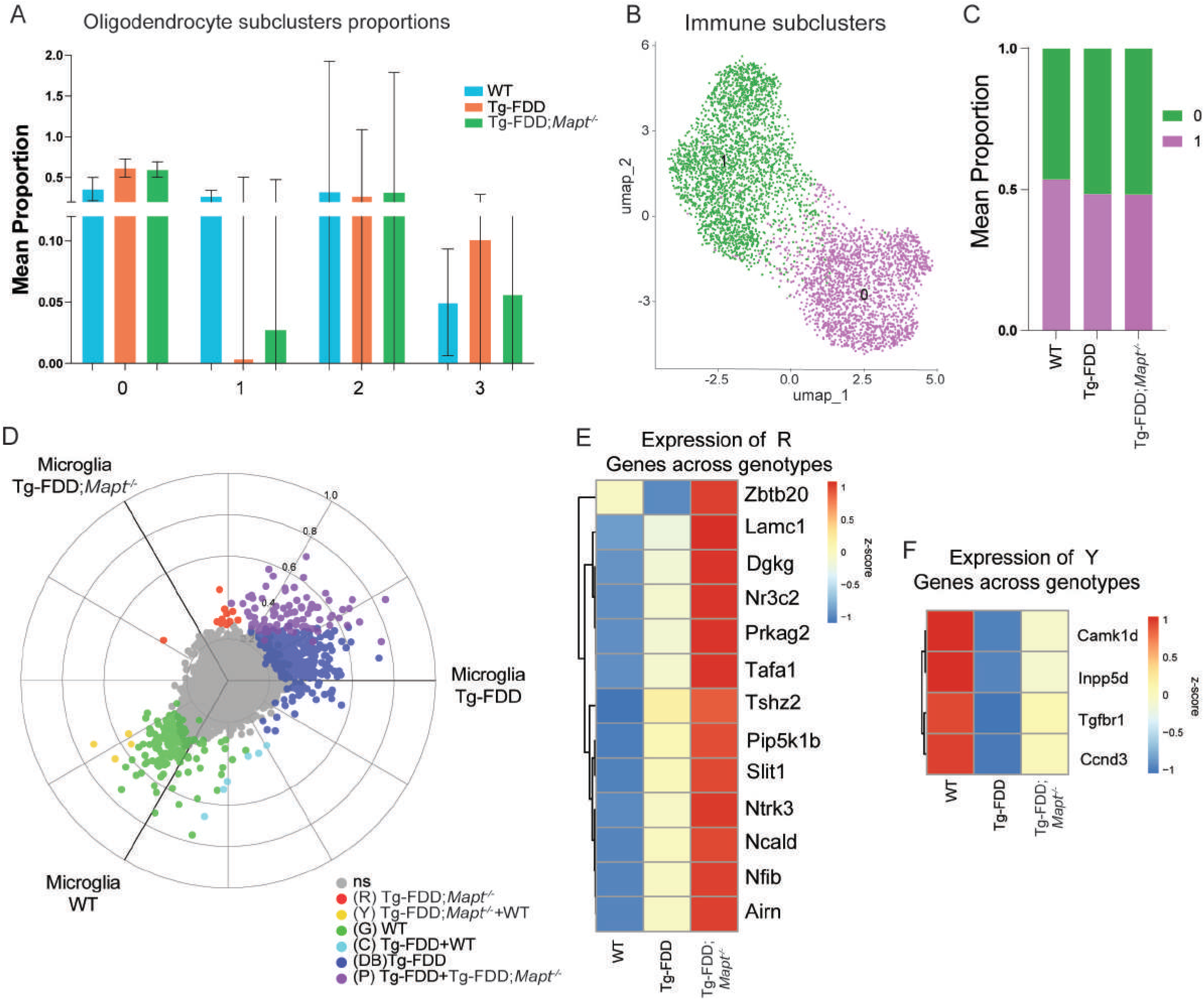
Oligodendrocyte proportion analysis and microglial transcriptional programs in Tg-FDD;*Mapt^-/-^* **A.** Barplot graph depicting cell-type proportion analysis comparing WT, Tg-FDD and Tg-FDD;*Mapt^-/-^*oligodendrocytes. Barplots data is shown as mean + SEM, one way ANOVA after logit-transformation**. B.** UMAP representation of immune cell subclusters. **C.** Stacked barplot graph depicting cell-type proportion analysis comparing WT, Tg-FDD and Tg-FDD;*Mapt^-/-^*immune cells**. D.** Microglia 3D Volcano plot showing DEGs for a three-way WT, Tg-FDD and Tg-FDD;*Mapt^-/-^* comparison. Each dot represents a gene. The X-, and Y- axes in the plot show the scaled mean expression differences for each of the three pairwise comparisons between genotypes. Genes further from the origin show larger differential expression. Dot color indicates statistical significance (one way ANOVA followed by BH multiple comparisons tests). Coordinates were calculated using the polar_coords() function from the volcano3D package. The radical scale axis shows normalized score scale. **E.** Heatmap depicting the Genes that belong to the microglia (R) set across genotypes. Row-scaled (z-scored). **F.** Heatmap depicting the Genes that belong to the microglia (Y) set across genotypes. Row-scaled (z-scored).

**Supplementary Figure 6.**
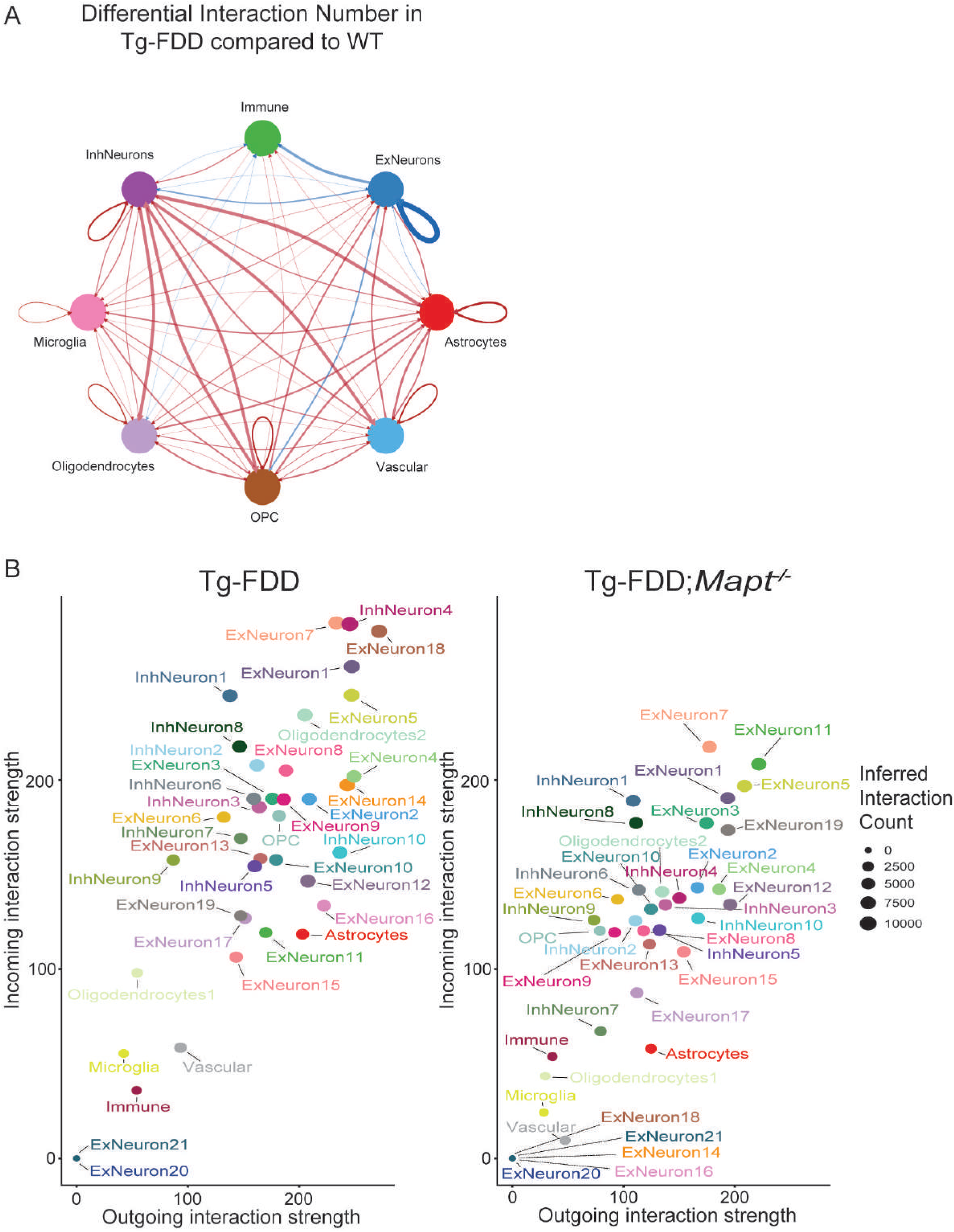
Predicted interaction strengths disease comparison of all cell clusters using CellChat. **A.** Chord plot showing the changes in interaction number between the main cell types in the brains of Tg-FDD mice compared with WT mice. Red lines indicate increased signaling, and blue lines indicate decreased signaling. The thickness of the lines represents the interaction strength. **B.** Predicted interaction strengths of all cell clusters in both conditions, computed from a merged CellChat object for WT (left) and *Mapt^⁻/⁻^* (right). **B.** Predicted interaction strengths of all cell clusters, computed from a merged CellChat object for Tg-FDD (left) and Tg-FDD;*Mapt^-/-^* (right). The x-axis shows the interaction strength of outgoing signals, and the y-axis shows the interaction strength of incoming signals.

